# Differences in size and number of embryonic type-II neuroblast lineages correlate with divergent timing of central complex development between beetle and fly

**DOI:** 10.1101/2024.05.03.592395

**Authors:** Simon Rethemeier, Sonja Fritzsche, Dominik Mühlen, Gregor Bucher, Vera S. Hunnekuhl

## Abstract

Despite its conserved basic structure, the morphology of the insect brain and the timing of its development underwent evolutionary adaptations. However, little is known about the developmental processes that create this diversity. The central complex is a brain centre required for multimodal information processing and an excellent model to understand neural development and divergence. It is produced in large parts by type-II neuroblasts, which produce intermediate progenitors, another type of cycling precursor, to increase their neural progeny. These neural stem cells are believed to be conserved among insects, but little is known on their molecular characteristics in insects other than flies. *Tribolium castaneum* has emerged as a valuable new insect model for brain development and evolution. However, type-II neuroblast lineages and their role in central complex development have so far not been studied in this beetle.

Using CRISPR-Cas9 we created a fluorescent enhancer trap marking expression of *Tribolium fez/earmuff*, a key marker for type-II neuroblast derived intermediate progenitors. Using combinatorial labelling of further markers including *Tc-pointed*, *Tc-deadpan*, *Tc-asense* and *Tc-prospero* we characterized the type-II neuroblast lineages present in the *Tribolium* embryo and their sub-cell-types. Intriguingly, we found 9 type-II neuroblast lineages per hemisphere in the *Tribolium* embryo while *Drosophila* produces only 8 per brain hemisphere. In addition, these lineages are significantly larger at the embryonic stage of *Tribolium* than they are in *Drosophila* and contain more intermediate progenitors. Finally, we mapped these lineages to the domains of early expressed head patterning genes. Notably, *Tc-otd* is absent from all type-II neuroblasts and intermediate progenitors, whereas *Tc-six3* marks an anterior subset of the type-II-lineages. The placodal marker *Tc-six4* specifically marks the territory where anterior medial type-II neuroblasts differentiate.

In conclusion, we identified a conserved pattern of gene expression in holometabolan central complex forming type-II neuroblast lineages, and conserved head patterning genes emerged as new candidates for conferring spatial identity to individual lineages. The higher number and greater lineage size of the embryonic type-II neuroblasts in the beetle correlate with a previously described embryonic phase of central complex formation which is not found in the fly. These findings stipulate further research on the causal link between timing of stem cell activity and temporal and structural differences in central complex development.

## Introduction

Neural development of insects allows to study molecular and cellular principles in easy to manipulate invertebrate models. The fruit fly *Drosophila melanogaster* (*Drosophila* hereafter) has mostly been used for this purpose, making it one of the best understood models for neurogenesis [1,2]. However, it has remained a puzzle how the huge ecological diversity of insects and the divergent neural anatomies that are adapted to different niches [3,4] evolved. Furthermore, *Drosophila* may in many instances not be a good representation of insect development and some processes are derived in the fly lineage [5–7]. For these reasons the beetle *Tribolium castaneum* (*Tribolium* hereafter) has been introduced as an additional model for insect neural development [8–11]. *Tribolium* is a grain storage pest and all stages live in flour, hence there is no major transition of habitat or lifestyle between the larval and adult stage as in flies. However, when compared to *Drosophila* the larvae are more mobile through the use of walking legs which allows them to navigate within or between food sources [3,6,10]. In this beetle many molecular genetic manipulation and labelling methods have been established [12–16]. Specifically, labelling of neural cell types in *Tribolium* has been facilitated by the advent of CRISPR-Cas9 [12,15,17]. Another factor that makes *Tribolium* an informative model for the understanding of insect brain development is the more conserved development of the head neuroectoderm facilitating the identification of specific neurogenic domains and allowing cross-species comparisons [11,18,19].

The insect central complex is an anterior, midline spanning neuropile and constitutes an important brain centre for the processing of sensory input, coordination of movement and navigation [20,21]. The highly conserved basic structure of the central complex, along with diverse specifications in response to ecological requirements among different insect species, make this brain structure a very interesting model for evolutionary developmental studies [3,10,20]. Between the two insect models *Drosophila* and *Tribolium* a temporal shift in the emergence of central complex neuropile was observed with *Tribolium* developing a functional larval central complex during embryogenesis whereas it develops only at the onset of adult life in *Drosophila* [3,10]. It is however not known how these temporal differences are established during development on a cellular and molecular level.

The entire insect central nervous system including the brain is produced by neural stem cells, the neuroblasts (NBs) [22]. While there are evolutionary modifications in the relative position of trunk NBs and their gene expression profiles between fly and beetle, the core determinants specifying their role as neural progenitors are conserved [8]. NBs undergo repeated divisions producing rows of ganglion mother cells (GMCs) which divide one more time to produce neurons and/or glia (see Fig. 1, top panel). This leads to neural cell lineages that include the neuroblast itself and its progeny [23,24]. Several NBs of the anterior-most part of the neuroectoderm contribute to the central complex, which has been characterized as the most complex brain structure (apart from the optic lobe) [25,26]. Intriguingly, in *Drosophila* and in the grasshopper *Schistocerca gregaria* a neuroblast subtype, the type-II neuroblasts (type-II NBs), were found to prominently contribute to the formation of the central complex [27,28]. These specific neuroblasts generate more offspring by producing another class of neural precursors, the intermediate progenitors (INPs), which also divide in a stem cell like fashion [29,30] (Fig. 1, lower panel). Type-II NBs and INPs have attracted a lot of attention because intermediate, cycling progenitors have also been described in vertebrates, the radial glia cells, which produce a variety of cell types [31,32]. In addition, in *Drosophila* brain tumours have been induced from type-II NB lineages [33], opening up the possibility of modelling tumorigenesis in an invertebrate brain, thus making these lineages one of the most intriguing stem cell model in invertebrates [34,35].

**Figure 1.**
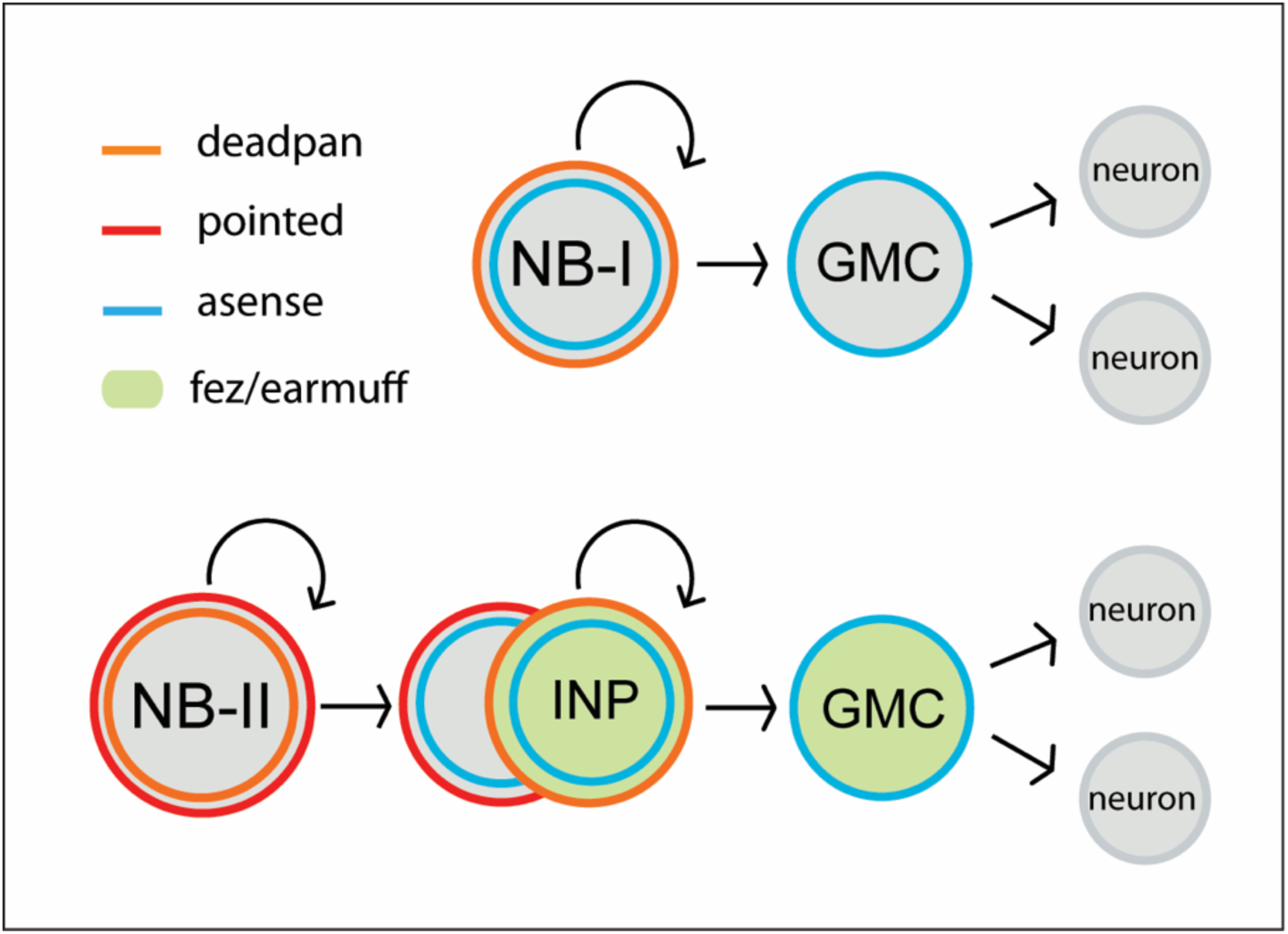
Neuroblasts of the *Drosophila* nervous system. Type-I neuroblasts (type-I NBs) (top panel) and type-II neuroblasts (type-II NBs) (lower panel). Type-I NBs undergo stem-cell-like divisions with each round producing a ganglion mother cell (GMC) that divides one more time to produce two neurons or glia cells. In *Drosophila*, all type-I NBs express *dpn* and *ase* whereas GMCs express only *ase*. Type-II NBs express *dpn* and *pnt* but not *ase*. They also undergo repeated divisions, with each round producing an intermediate progenitor cell (INP) which also is a proliferating progenitor and expresses *ase*. INPs undergo a maturation process during which *pnt* is still expressed initially, but not *dpn*. Mature INPs express *dpn*, *ase* and *fez/erm* whereas GMCs express *ase* and *fez/erm*. Type-II NB lineages are only found in the anterior brain, they produce central complex neurons and glia [30,36].

In the developing insect brain the number of neurons that is produced by each type-II NB is significantly larger than the offspring of a type-I NB. Medial *Drosophila* type-II NBs produce an average of 450 neurons whereas type-I NBs of the brain only generate 100-200 neurons each [37–39]. This increased number of neural cells that are produced by individual type-II lineages along with the fact that neuroblast lineages can produce different types of neurons, leads to the generation of extensive neural complexity within the anterior insect brain [27,30].

Based on descriptions of large proliferative lineages in the grasshopper *Schistocerca gregaria,* a hemimetabolous insect distantly related to flies, it is widely believed that type-II NBs and INPs are conserved within insects [25,40]. However, a molecular characterisation of such lineages that on the one hand will allow to test whether the cell type markers characterized in *Drosophila* are more widely conserved, and on the other hand will facilitate an identification of sub-cell types, has not been performed in an insect other than *Drosophila*.

Whereas it is widely believed that *Tribolium* does use type-II NB in central complex formation [3,17,41] their presence has so far not been unequivocally shown and a thorough comparison of type-II NB lineages and their sub-cell-types between the fly and the beetle model has been lacking.

The characterization of type-II NBs in *Drosophila* has in large parts been based on a specific reporter line in which eGFP expression is driven by an *earmuff (fez/erm)* enhancer element and marks INP-lineages [30,42,43]. *Tribolium earmuff* has been first described as *Tc-fez*, referring to the vertebrate *earmuff* ortholog *fez* [19]. We will use *Tc-fez/erm* in the following. A defined sequence of gene expression has been described in *Drosophila* type-II NBs, INPs and GMCs. The ETS-transcription factor *pointed* (*pnt*) marks type-II NBs [44,45], which do not express the type-I NB marker *asense* (*ase*) but the neural gene deadpan (*dpn*) (Fig. 1). Intermediate progenitors express *fez/erm* and *ase* and at a more mature stage also *dpn*, whereas *prospero* (*pros*) marks mature INPs and GMCs [30,46]. *Six4* is another marker for *Drosophila* type-II NBs and INPs [47].

In the present work we characterize type-II neuroblasts in *Tribolium* with respect to their number, their location and the conservation of the molecular markers known from *Drosophila*. We also wanted to test, in how far the earlier emergence of the central complex in beetles would be reflected by a change of division activity of type-II NBs or intermediate progenitors. To that end, we used CRISPR-Cas9 induced genome editing for creating a *Tc-fez/erm* enhancer trap line to characterize embryonic *Tc-fez/erm* -expressing cells including INPs and GMCs of type-II NB lineages and combined this line with other labelling methods including multicolour *hybridisation chain reaction* (HCR) [48,49]. We found that the *Tribolium* embryo produces 9 *Tc-pnt*-expressing type-II NBs compared to only 8 type-II NB lineages in the *Drosophila* embryo [50]. We show that these lineages produce central complex cells and confirm a largely conserved molecular code that differentiates INPs of distinct maturation stages and GMCs. Intriguingly, we found that lineage sizes in *Tribolium* embryos are considerably larger and include more mature INPs than in the *Drosophila* embryo [50]. We also show that the placodal marker gene *Tc-six4* characterizes the embryonic tissue that gives rise to the larger anterior group of type-II NBs, whereas *Tc-six3* is marking only an anterior subset of these lineages. Interestingly there is a part of the central complex precursors expressing neither *Tc-six3* nor *Tc-otd,* which is absent from all type-II NB lineages.

## Results

### A CRISPR-Cas9-NHEJ generated enhancer trap lines marks *Tc-fez/erm*-expressing cells at the embryonic and larval stage

We created an enhancer trap line driving eGFP that reflects *Tc-fez/erm* gene expression in the embryo. We named the line *fez magic mushrooms* (fez-mm-eGFP) because it marks the mushroom bodies of the larval brain (Fig. 2 A-C). The reporter construct was inserted 160 bp upstream of the *fez/erm*-transcription start site using the *non-homologous end joining* (NHEJ) repair mechanism (see supplementary Fig. S1 for scheme). Enhancer traps sometimes only reflect parts of a gene expression pattern, therefore we carefully evaluated eGFP co-expression with *Tc-fez/erm*-RNA. We found that at the embryonic stage fez-mm-eGFP co-localized with most *Tc-fez/erm*-expressing cells in an area that will give rise to the protocerebral structures mushroom bodies, larval eyes and central complex [11,19] (Fig. 2, panel D). The fez-mm-eGFP line marked several progenitor cell types including cells that we characterize as type-II NB-derived intermediate progenitors (see below). Based on these analyses, we use the fez-mm-eGFP expression as a reporter for *Tc-fez/erm* expression.

**Figure 2.**
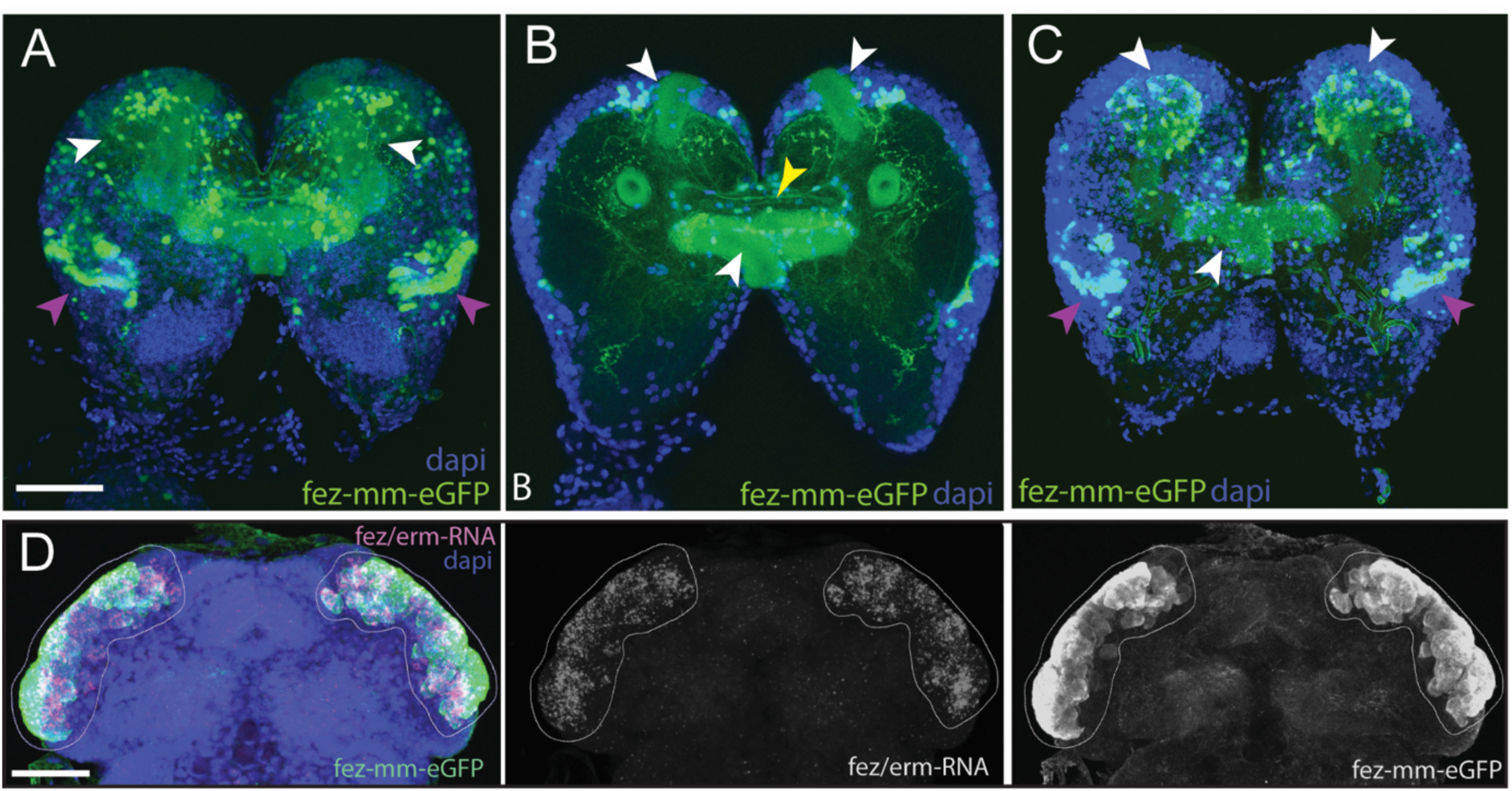
Magic mushrooms (fez-mm-eGFP) reporter line. A-C) eGFP-expression in a larval instar brain, D) EGFP antibody labelling in combination with *Tc-fez/erm* RNA *in situ* in the embryonic head, A-D) Dapi labelling of nuclei. Scalebars in A and D are 50 µm. **A)** Full projection of whole brain scan (4^th^ instar larvae): fez-mm-eGPF expression marks many cells of the mushroom bodies (white arrowheads) and of the larval optic lobe (purple arrowheads). Note that we use the terms dorsal and ventral with respect to the neuraxis of the larva. See [10] for orientation. **B)** Central part of the whole brain scan (projection of substack, same brain as in A): mm-eGFP marks the peduncles of the mushroom bodies (white arrowheads) and some central complex ensheathing cells (probably glia cells; yellow arrowhead). **C)** Projection of the dorsal part of same brain as in A shows calyces and peduncles of mushroom bodies (white arrowheads), and optic neuropile (purple arrowheads). **Panel D)** Extensive co-localization of fez-mm-eGFP with *Tc-fez/erm*-RNA in white encircled area which gives rise to protocerebral structures. Projection of all planes where expression was detected, embryonic stage NS11.

### *Tc-pointed* marks a population of nine type-II NBs that are associated with fez-mm-eGFP marked INP-lineages

To identify type-II NBs, we searched for cell groups that express the markers known from flies. The transcription factor *pointed* (*pnt*) marks type-II NBs in *Drosophila* and is required for their correct differentiation [44,45]. In both *Tribolium* and *Drosophila*, *pnt* is also expressed in other areas (this work/[51,52]). Therefore, we looked for *Tc*-*pnt* expression in large cells with large nuclei (typical NB morphology) that were closely associated with *Tc-fez/erm*-expressing cells (marking putative INPs). We found a total of 9 such clusters instead of the 8 expected from flies (Fig. 3 A-C). We also found that the large *Tc-pnt* expressing cells as well as fez-mm-eGFP expressing cells are mitotically active (Fig. 3 C-I). Further molecular and cell size analysis corroborated our interpretation that these cells constitute type-II NBs, intermediate progenitors (INPs) at different maturation stages and ganglion mother cells (GMCs) (see details below).

**Figure 3.**
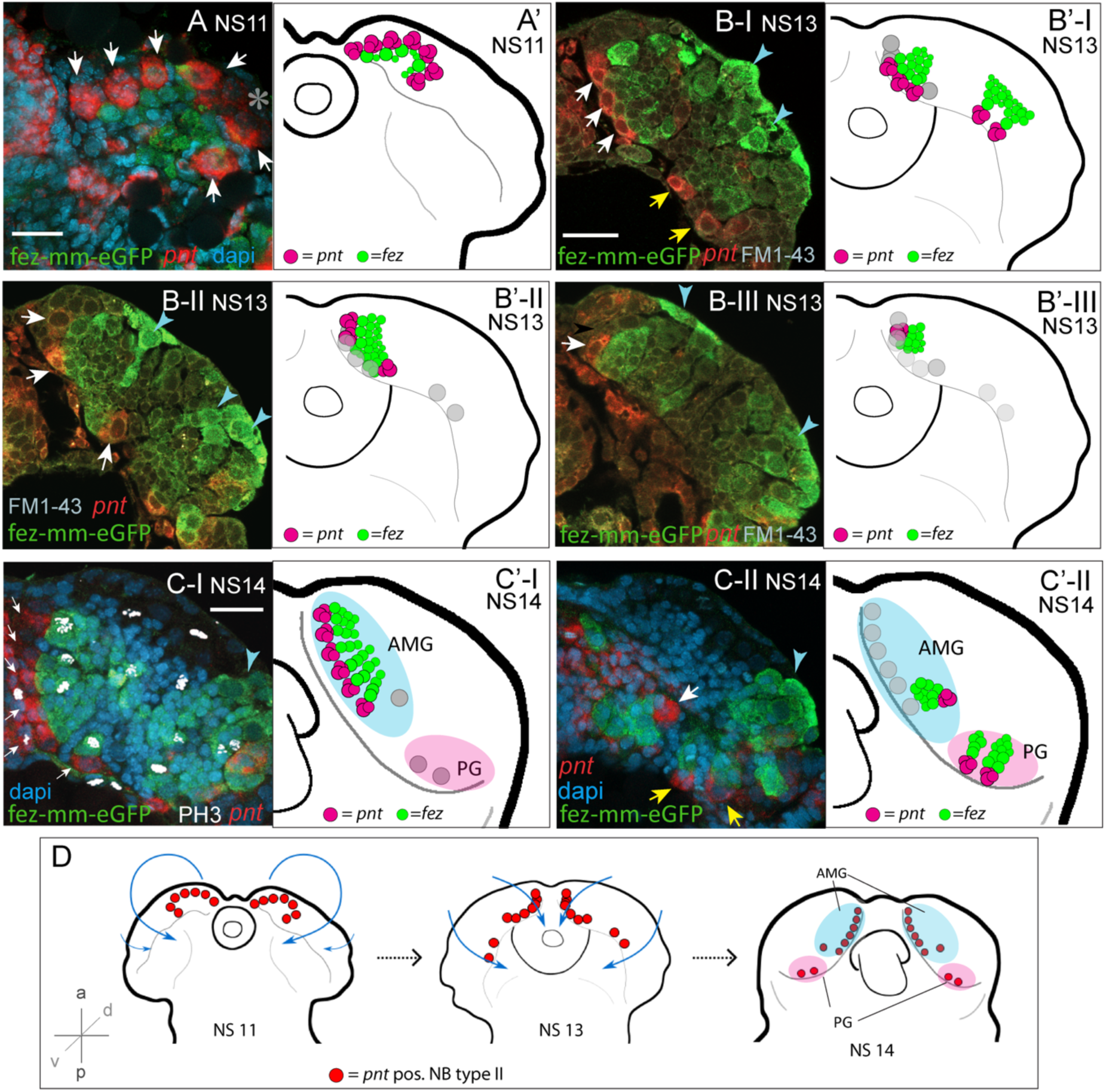
Developmental occurrence and progression of *Tc-pnt*-clusters and associated fez-mm-eGFP cells (type-II NB lineages). A-C) Anti-GFP antibody staining reflecting fez-mm-eGFP expression in combination with *Tc-pnt* RNA *in situ* hybridisation and dapi labelling of nuclei (A, C) or FM1-43 membrane staining (B), single planes from confocal z-stacks of right head lobes. Scalebars are 25 µm. A’-C’) Schematic drawing of the head lobe at the respective stages; clusters out of focus depicted by grey circles. D) Schematic depiction of the localization of type-II NBs during *Tribolium* head morphogenesis. **A)** At stage NS11 seven *Tc-pnt*-clusters (white arrows) are found in an anterior-medial horseshoe-like arrangement with fez-mm-eGFP cells in the centre. Asterisk indicates the position of one *Tc-pnt*-cluster which is outside the focal plane of this image. **B)** At stage NS13 the seven anterior-medial clusters have been moved to the medial margin of the developing brain by morphogenetic movements of the head lobes (white arrows). The clusters are situated in different dorso-ventral levels shown in the different images B-I to B-III. Two additional posterior *Tc-pnt*-clusters have appeared (yellow arrows). Blue arrows point to fez-mm-eGFP positive cells that are not part of the type-II NB lineages. **B-I**: ventral level **B-II** mid-level, **B-III**: dorsal level. **C)** At stage NS14 a group of six clusters are arranged in one plane anterior-medially in the brain lobe (white arrows) (**C-I).** Anti-PH3 labelling marks mitotic cells within the lineages (white signal) and clusters out of the focal plane are indicated by grey circles in C’-I. **C-II** One cluster of the anterior median group (white arrow) is in a deeper plane, as well as the two posterior clusters (yellow arrows). Blue arrowheads point to lateral fez-mm-eGFP expression not associated with type-II NBs. A’-C’) AMG =anterior medial group (blue), PG =posterior group (pink). Grey circles in B’-C’ indicate position of type-II NBs in different focal planes. **D)** *Tc-pnt*-expressing cells identified as type-II NBs first appear bilaterally in the anterior most part of the embryonic head (stage NS11). During head morphogenesis this anterior tissue folds over so that the type-II NBs end up in a more medial position. Two more type-II NBs emerge posteriorly (stage 13). At the end of embryogenesis (stage NS14) the type-II NBs can be divided in an anterior medial group (AMG (blue); including type-II NBs 1-7, numbering starting from anterior) and a posterior group (PG (pink), including type-II NBs 8-9).

Looking at a series of embryonic stages we determined the emergence and further development of *Tc-pnt* and fez-mm-eGFP expressing lineages as well as their arrangement with respect to the embryonic head and to one another. We first detected conspicuous adjacent expression of *Tc-pnt* and fez-mm-eGFP at stage NS7 (not shown). At stage NS11 seven *Tc-pnt*-clusters are present in a horse-shoe-shaped formation surrounding fez-mm-eGFP positive cells (white arrows in Fig. 3 A, star marks one cluster out of focus). At stage NS13 these groups are found in a position at the medial border of the head lobes and their number has increased to nine. The seven clusters that were already present at stage NS11 are arranged in an anterior group (white arrowheads in Fig. 3B) at stage NS13 while two additional clusters have emerged more posteriorly (yellow arrows in Fig. 3B-I). The clusters are in different depths (reflected by the three optical sections shown in Fig. 3 B-I to B-III, clusters out of focus are visualized by grey circles in Fig. 3 B’I-II, respectively). Fez-mm-eGFP expression not related to type-II NB offspring was found in the lateral head lobe from NS13 onwards (blue arrowheads in Fig. 3). At stage NS14 six of the anterior-medial clusters are aligned in one plane along the medial rim of the developing brain (Fig. 3 C-I), whereas the most posterior cluster of the anterior group is located at a separate position and in a deeper plane (Fig. 3C-II). The posterior group is found at about the same focal plane as well (Fig. 3 C-II). The aligned six clusters produce their *Tc-fez/erm*-positive offspring towards laterally whereas the *Tc-fez/erm*-positive offspring of the more posterior cluster is found medially of it (Fig. 3 C-II). The two most posterior clusters produce *Tc-fez/erm*-cells in an anterior-lateral direction (Fig. 3 C-II). The apparent re-arrangement of the *Tc-pnt*-clusters from the anterior rim of the embryonic head to a medial position reflects not active migration but follows the overall tissue movements during head morphogenesis that was previously described in *Tribolium* [19] (Fig. 3 D). Based on their position by the end of embryogenesis (stage NS14) we assign the nine type-II NB clusters into two groups: one anterior median group consisting of seven type-II NBs and one posterior group consisting of two type-II NBs (see Fig. 3 C and D).

### Characterization of cell- and nuclear size of *pnt*-positive type II NBs

We wanted to confirm that each of the *Tc-pnt* and *Tc-fez/erm* expressing lineages contains a neuroblast. Neuroblasts, including type-II NBs, are larger than other cells, have larger nuclei and are mitotically active [40]. Therefore, we determined the cell size of the largest cells of the *Tc-pnt* cluster and compared it to a random sample of cells of the embryonic head. Both samples were taken from stage NS13 and NS14 embryos. We found that the cells that we had assigned as Type-II NBs had a significantly larger diameter (av.: 12.98 µm; n=43) than cells of the reference sample (av.: 6.75 µm; n=1579) (Fig. 4). We also found that the diameters of nuclei of type-II NBs were significantly larger (av.: 8.73 µm; n=17) than the ones of a control sample (av.: 5.21 µm; n=1080) (Fig. 4). Although type-II NBs were clearly larger than the average cell of the head lobes the control sample also contained cells of an equal size (figure 4) which may constitute type-I NBs.

**Figure 4.**
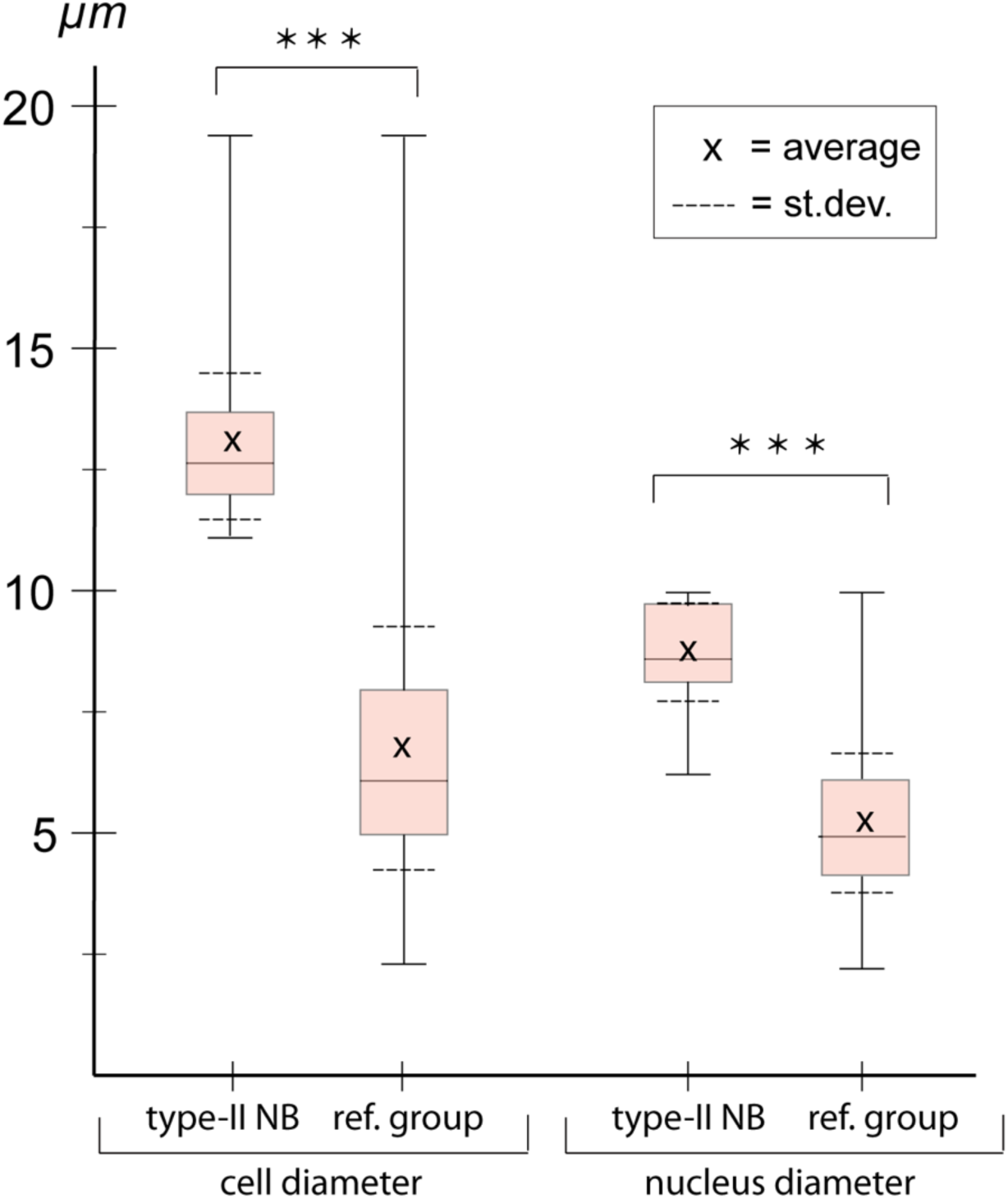
Cell- and nuclear diameter of type-II NBs compared to a control group. Stages NS13-14. Left: diameter of *Tc-pnt*-pos. type-II NBs (average diameter=12.98 µm; n=43) compared to control group (average diameter=6.75, n=1579). Right: average nuclear diameter of type-II NBs (=8,73 µm, n=17) compared to average nuclear diameter of control group (=5.21, n=1080). Differences between groups are significant (t-test; *** ≘ P < 0.001).

### Conserved patterns of gene expression mark *Tribolium* type-II NBs, different stages of INPs and GMCs

*Drosophila* type-II NBs, INPs and GMCs express a specific sequence of genes including *asense* (*ase*), *deadpan* (*dpn*) and *prospero* (*pros*) (Fig. 1) [30]. However, these markers have up to now not been tested for expression in type-II lineages in other organisms. We tested if the respective lineages express these markers in a comparable sequence in the beetle and if they mark distinctive cell types within the lineages (Fig. 5 panel A-D and Fig. panel 6 A, B). We found that the large *Tc-pnt*-positive but fez-mm-eGFP negative cells (i.e. type-II NBs) express the gene *Tc-dpn*, in line with expression of this gene in *Drosophila* neural precursors (Fig. 5, panel A). Like *Drosophila* type-II NBs, these cells do not express the type-I NB marker *Tc-ase* (yellow arrowhead in Fig. 5, panel D). We further found additional cells directly adjacent of the type-II NBs itself which we believe are recently born immature INPs. They are *Tc-pnt*-positive but fez-mm-eGFP-negative and neither express *Tc-ase* (Fig. panel D, pink arrowheads). Expression of *Tc-ase* starts in the following in immature *Tc-pnt*-positive INPs (Fig. 5 panel D; blue arrowhead) and is maintained in INPs that are *Tc-pnt*-negative but are marked by fez-mm-eGFP (Fig. 5, panel D and E). *Tc-dpn* is absent from immature *Tc*-*pnt*-positive INPs (Fig. 5D, pink arrowhead) but is expressed in mature INPs (orange arrowheads in Fig. 5, panel B). Cells that are located at the distal end of the lineages that do not express *Tc*-*dpn* but are positive for fez-mm*-*eGFP and *Tc*-*ase* are classified as GMCs (Fig. 5, panels B, C and E). We found the gene *Tc-pros* expressed in most fez-mm-eGFP expressing cells of a lineage (in Fig. 6, panel A-B) and infer from the extent of the expression that it is expressed in mature INPs (*Tc-dpn*+) and GMCs (*Tc-dpn-*). Fez-mm-eGFP positive cells at the base of the lineage that do not express *Tc-pros* are immature INPs (Fig. 6, panel A, B; blue arrowheads). In summary, the expression dynamics found in the *Tribolium* type-II NB lineages are very similar to the one found in *Drosophila.* Therefore these neural markers can be used for a classification of cell types within the lineages into type-II NBs (*Tc-pnt*+, *Tc-fez/erm*-, *Tc-ase*-, *Tc-dpn*+, *Tc-pros*-), immature-I INPs (*Tc*-*pnt*+, *Tc-fez/erm*-, *Tc-ase-, Tc-dpn-, Tc-pros-*), immature-II INPs (*Tc-pnt*+, *Tc-fez/erm*+, *Tc-ase*+, *Tc-dpn*-, *Tc-pros*-), mature INPs (*Tc-pnt*-, *Tc-fez/erm*+, *Tc-ase*+, *Tc-dpn*+, *Tc-pros*+), and GMCs (*Tc-pnt*-, *Tc-fez/erm*+, *Tc-ase*+, *Tc-dpn-, Tc-pros*+). This classification is summarized in Fig. 7 A-B. Image stacks from specimens co-stained with the anti-PH3 mitosis marker (data shown in fig. 3C and listed in table 1 and 2, available at https://doi.org/10.25625/8IVICL) indicate that immature-I and -II INPs are not dividing whereas mature INPs are.

**Figure 5:**
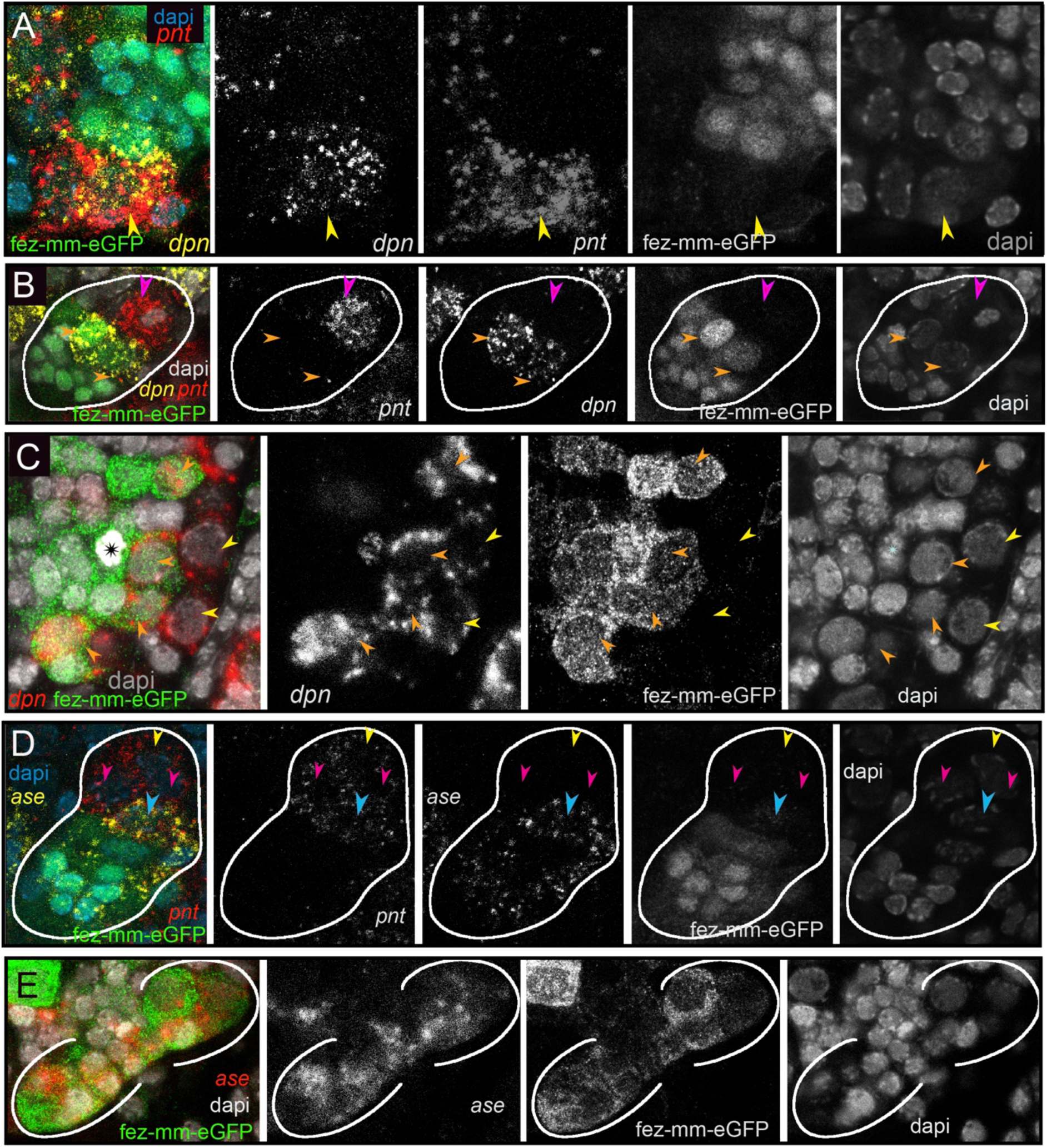
Differential expression of the neural markers *Tc-pnt, Tc-fez/erm, Tc-dpn* and *Tc-ase* in type-II NBs, INPs and GMCs. A, B, D) Anti-GFP staining visualizing fez-mm-eGFP expression in combination with HCR-labelling of further factors. C, E) GFP antibody staining in combination with RNA *in situ* hybridisation. A-E) Dapi staining of nuclei. All panels are single planes from confocal z-stacks. **Panel A)** Stage NS12, expression of *Tc-pnt* and *Tc-dpn* in type-II NBs (yellow arrowhead) with adjacent fez-mm-eGFP positive INPs. **Panel B)** Stage NS13, white line marks one lineage. Expression of *Tc-dpn* is absent in the immature-I *Tc-pnt*+ INPs (pink arrowhead) but present in mature INPs (orange arrowheads), white line marks one lineage. Type-II NB not in focus of this image. **Panel C)** Stage NS14, expression of *Tc-dpn* in type-II NBs (yellow arrowheads), and mature INPs (orange arrowheads). Asterisk marks mitosis in a fez-mm-eGFP cell visible through the condensation of chromatin stained by dapi. **Panel D)** Stage NS13, *Tc-ase*- and *Tc-pnt*-expression in type-II NBs and INPs, white line marks one lineage. Yellow arrowhead: type-II NBs (*Tc-pnt*+), pink arrowheads: immature-I INP (*Tc-pnt*+), blue arrowhead: immature-II INP (*Tc-fez*+/ *Tc-ase*+/ *Tc-pnt*+). **Panel E)** Stage NS13, expression of *Tc-ase* in fez-mm-eGFP positive cells, location of type-II NBs and immature-I INPs is outside the focus.

**Figure 6.**
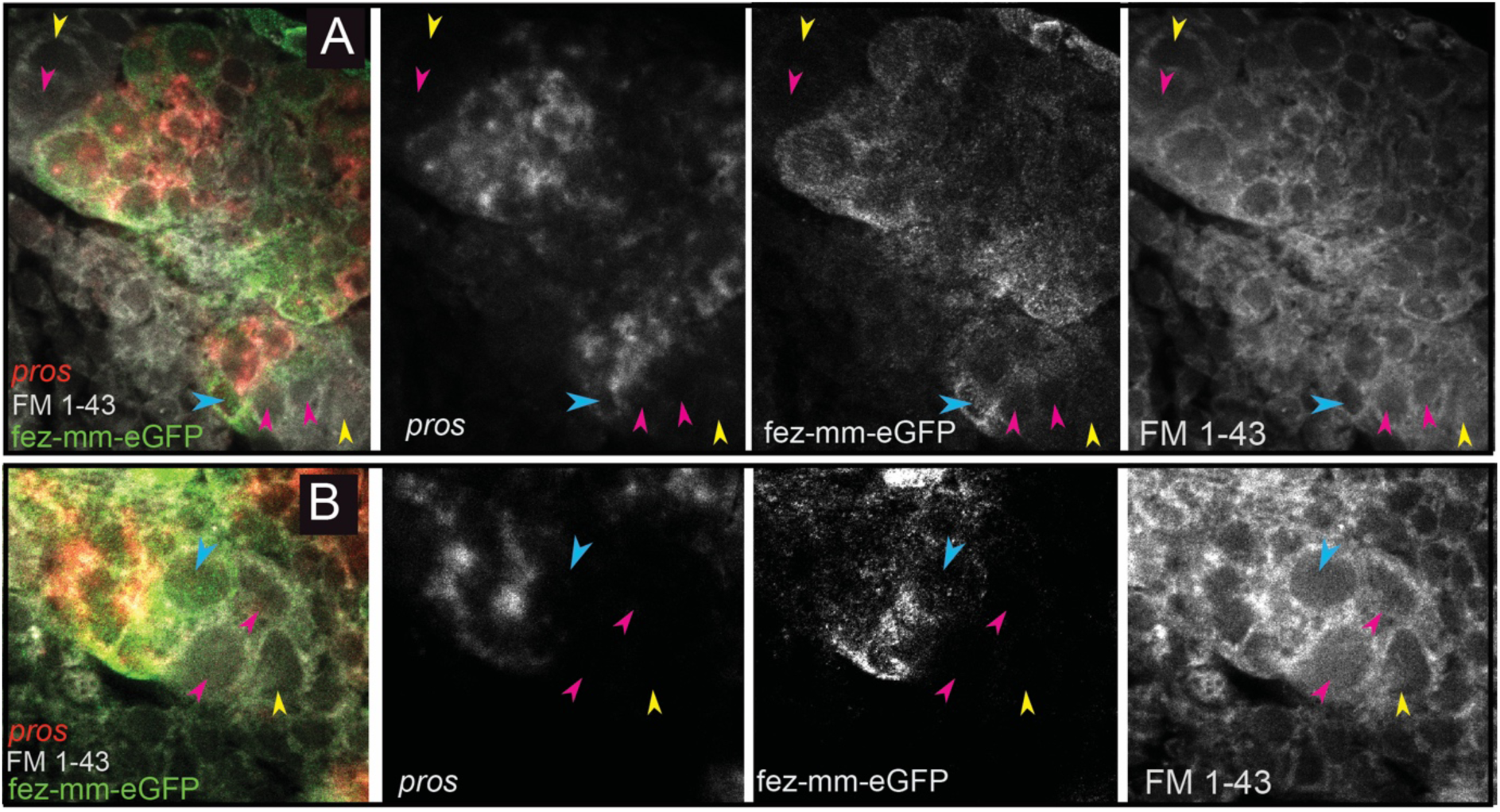
Expression of *Tc-pros* in fez-mm-eGFP cells. A-B) Anti-GFP antibody staining in combination with *Tc-pros* RNA *in situ* hybridization. Membranes stained with FM 1-43. Single planes from confocal z-stacks. **Panel A)** Stage NS13, two type-II NB lineages producing INPs and GMCs in opposing directions. Expression of *Tc-pros* in mature INPs and GMCs. Cluster of type-II NBs (yellow arrowheads) and immature-I INPs (pink arrowheads). Immature-II INPs express *Tc-fez/erm* but not *Tc-pros* (blue arrowhead). **Panel B)** Stage NS13, detail of *Tc-pros*-negative type-II NBs (yellow arrowhead) and immature-I INPs (pink arrowheads) and immature-II INP (blue arrowhead). Note that distinction between type-II NB and immature INPs in **A)** and **B)** is only made based on position.

**Figure 7.**
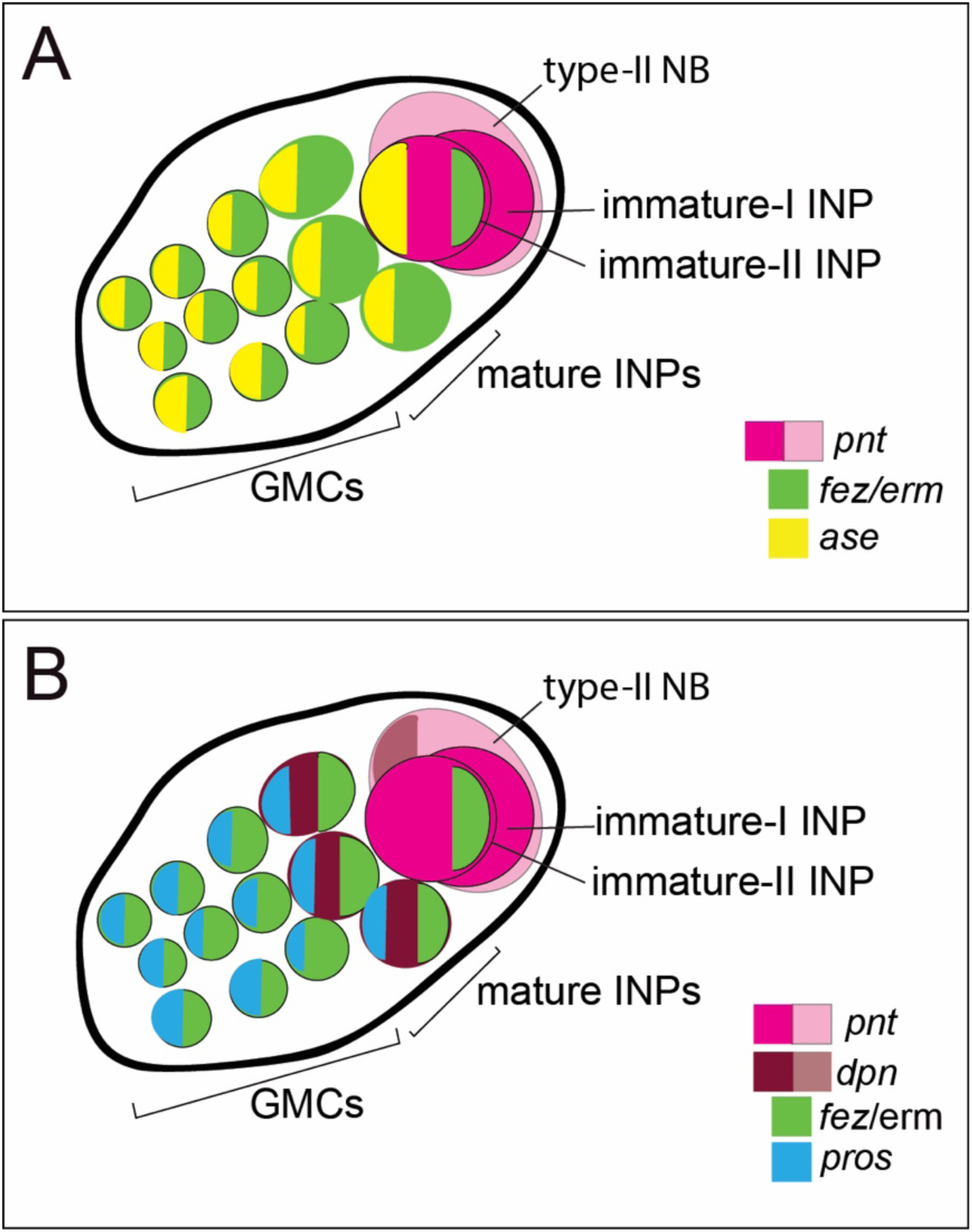
Schematic drawing of expression of different markers in a type-II NB lineage. A) and B) show the same lineage with *Tc-fez/erm* and *Tc-pnt* expression but with different additional markers mapped on top. **A)** Subclassification into type-II NBs (*Tc-pnt+*), immature-I INPs (*Tc-pnt+*), immature-II INPs (*Tc-pnt+, Tc-fez/erm+, Tc-ase+*) and mature INPs and GMCs (*Tc-fez/erm+, Tc-ase+*) (based on stainings shown in Fig. 5). **B)** Type-II NB (*Tc-pnt+, Tc-dpn+*), immature-I (*Tc-pnt+*) and immature-II (Tc-pnt+, *Tc-fez/erm+*). Mature INPs express *Tc-dpn*, *Tc-pros* and *Tc-fez/erm* whereas GMCs express only *Tc-fez/erm* and *Tc-pros* (based on staining shown in Fig. 5 and 6). Note that of the selected markers immature-I INPs express *Tc-pnt* only.

**Table 1:** *Tc-fez/erm* positive cell number (type-II NBs, INPs and GMCs)

| Embryo#/stage | Total number of cells of 7 anterior lineages | % dividing cells (PH3 antibody) | Cells per lineage (average) |
| --- | --- | --- | --- |
| 1/NS 13 | 146 | 8.2% | 20.9 |
| 2/NS 13 | 111 | 6.3% | 15.9 |
| 3/NS 14 | 139 | 5.0% | 19.9 |
| 4/NS 14 | 96 | 10.5% | 13.7 |

### Type-II NB lineages produce central complex cells that are marked by shaking hands (skh)

To test if the *Tribolium* embryonic type-II NB lineages contribute to the beetle central complex like in the fly, we used the shaking hands*-* (skh-) reporter line that marks central complex neurons and their postmitotic embryonic precursors [41]. By co-staining for *Tc-pnt* and *Tc-fez/erm* RNA in the background of the skh-line we found that the skh-positive cells are located directly distally of the type-II NB lineages in a position where the type-II NBs derived neural cells are expected (Fig. 8 panels A-B, E,F). Most skh-cells have switched off *fez/erm*-expression, but at the transition between *Tc-fez/erm* and skh cells we found some cells that express both markers (Fig. 8 panel D, E), supporting the view that many skh-positive central complex cells stem from the *fez*-expressing GMCs of the type-II NB lineages (Fig. 8 E). We found skh cells at the end of lineages of the anterior-median group (Fig. 8, panel A) and at the end the two posterior lineages (Fig. 8, panel B). We therefore conclude that both groups contribute to central complex neuropile (Fig. 8 F). We cannot say whether the lineage of type-II NB seven, which we assigned to the anterior-median group, but which is located with some distance to the other type-II NB of that group (Fig. 3 C, D; grey circle in Fig. 8 F), produces skh+ cells, but we assume that it does.

**Figure 8.**
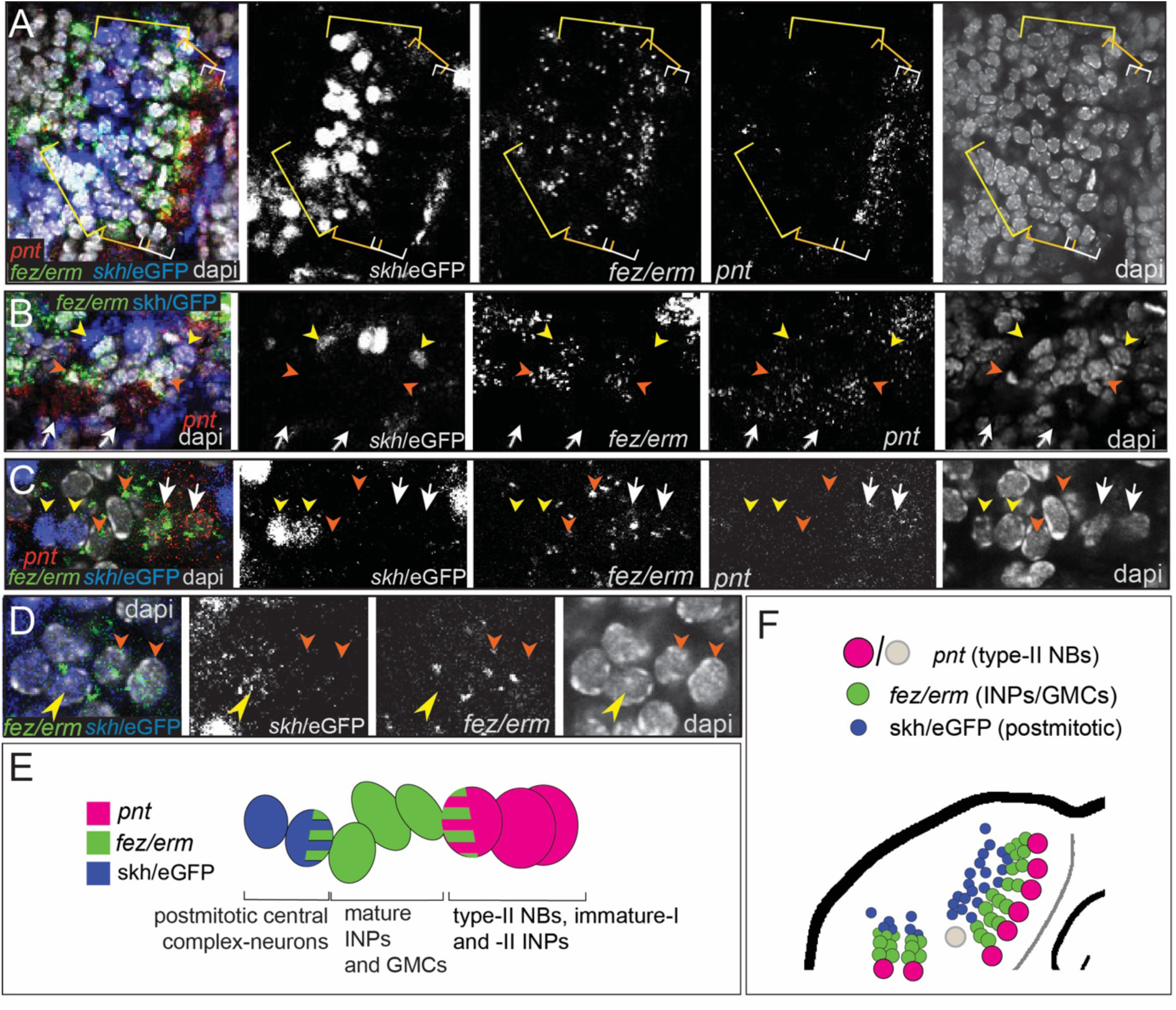
Expression of the transgenic central complex reporter skh/eGFP in relation to type-II NB lineages. A-D) Antibody labelling of eGFP, which is expressed in the pattern of skh [41] in combination with HCR visualizing *Tc-fez/erm* and *Tc-pnt*, dapi labelling of nuclei, single planes of confocal z-stacks, all stage NS14. **A)** Left head lobe, anterior group of *Tc-pnt*-positive type-II NBs, *Tc-fez/erm*-expressing INPs/GMCs and skh/eGFP positive postmitotic cells. **B)** Left head lobe, posterior group of type-II NB lineages. *Tc-pnt+* type-II NBs or immature-I/-II INPs (white arrows), *Tc-fez/erm*+ INPs/GMCs (orange arrowheads) and skh/eGFPpostmitotic cells (yellow arrowheads). **C)** Detail of one lineage including type-II NB cluster (*Tc-pnt*+, white arrows), INPs/GMCs (*fez+,* orange arrowheads) and skh/eGFPcells at the end of the lineage (yellow arrowheads). **D)** Detail of transition from *Tc-fez/erm*+ cells (orange arrowheads) to skh/eGFPcells reveals a small area of co-expression of both factors (yellow arrowhead). **E)** Schematic drawing of type-II NB lineage showing relative positions and markers of type-II NBs, INPs/GMCs and postmitotic central complex forming cells. **F)** Schematic overview of the arrangement of the type-II NB lineages including skh/eGFP+ central complex forming cells in the head lobe of stage NS14 (immature-I/-II INPs not shown for simplicity). No data on skh-GFP expression in lineage of type-II NB 7 (marked in grey) is available.

### The *Tribolium* embryonic lineages of type-II NBs are larger and contain more mature INPs than those of *Drosophila*

In beetles, a single-unit central body (most likely constituting the fan-shaped body [10]) develops during embryogenesis and has gained functionality at the onset of larval life [15]. By contrast, in *Drosophila* central complex commissural tracts originating from type-II NBs form in the embryo but the type-II NB become quiescent and only resume divisions in the late larval stage [29,50]. In stark contrast to *Tribolium* a functional central complex is only observed in the adult fly [3]. In addition, the volume of the L1 central brain including the central complex is about four times larger in *Tribolium* than it is in *Drosophila* [53], while there are only small differences in the relative neuropilar volumes of the adult beetle and fly central bodies [53,54]. As type-II NBs contribute to central complex development we asked in how far the embryonic division pattern of these lineages in the beetle would reflect this heterochronic development. To assess the size of the embryonic type-II NB lineages in beetles we counted the *Tc-fez/erm* positive (fez-mm-eGFP) cells (INPs and GMCs) of the anterior medial group (type-II NB lineages 1-7). As we were not always able to distinguish between cells belonging to neighbouring INP-lineages we quantified and averaged the cell number of all lineages of the anterior medial group. We also evaluated the proportion of dividing cells in that group based on anti-PH3 staining (table 1).

We found that at stage NS13 (early retracting embryo) each lineage consisted on average of 18,4 progenitor cells and at stage NS14 (retracted embryo) of 16.8 progenitors. Between 5-10 % of the cells were marked by the PH3 antibody. As PH3 marks dividing cells in a specific phase, the portion of dividing cells may be higher. We further quantified the number of mature *Tc-dpn*-positive INPs (table 2).

**Table 2:**
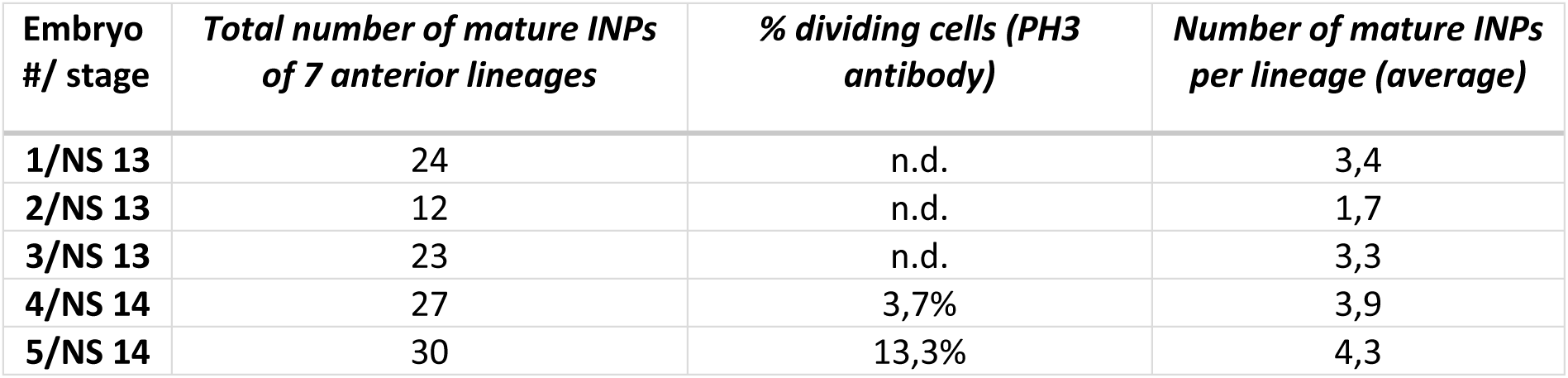
Mature INPs (*Tc-fez/erm* and *Tc-dpn* positive)

| Embryo #/ stage | Total number of mature INPs of 7 anterior lineages | % dividing cells (PH3 antibody) | Number of mature INPs per lineage (average) |
| --- | --- | --- | --- |
| 1/NS 13 | 24 | n.d. | 3,4 |
| 2/NS 13 | 12 | n.d. | 1,7 |
| 3/NS 13 | 23 | n.d. | 3,3 |
| 4/NS 14 | 27 | 3,7% | 3,9 |
| 5/NS 14 | 30 | 13,3% | 4,3 |

We used the numbers from *Tribolium* (table 1, 2) for comparison with type-II NB lineages from *Drosophila* embryos (data from [29]). At fly embryonic stage 13, which is comparable to *Tribolium* stage NS14 in that is represents a retracted germband, lineages only contained one or more INPs but no GMCs [29]. Therefore, we compared our data for the anterior lineages of *Tribolium* stages NS13 and NS14 to the anterior and median cluster in the *Drosophila* stage 15 and 16 embryos. In both species these stages are the two penultimate stages of embryogenesis when all type-II NB lineages are present, and importantly, in *Drosophila* the maximum lineage sizes and numbers of INPs are reached at these stages. We found that the beetle lineages were significantly larger than the corresponding ones in *Drosophila* (Fig. 9; *Drosophila* data taken from [29]). Of note, cell counts in *Drosophila* also included neurons [29]. Our counts are based on *Tc-pnt* and fez-mm-eGFP expression and therefore mostly included progenitors. However, based on the small area of co-expression of *Tc-fez/erm* and skh-expression (see above/ figure 8) they may also include some newly born neurons, but the counts in *Drosophila* included more mature neurons which further increases the difference in the number of progenitor cells between the two species and the observed difference in lineage size represents a minimum estimate.

**Figure 9.**
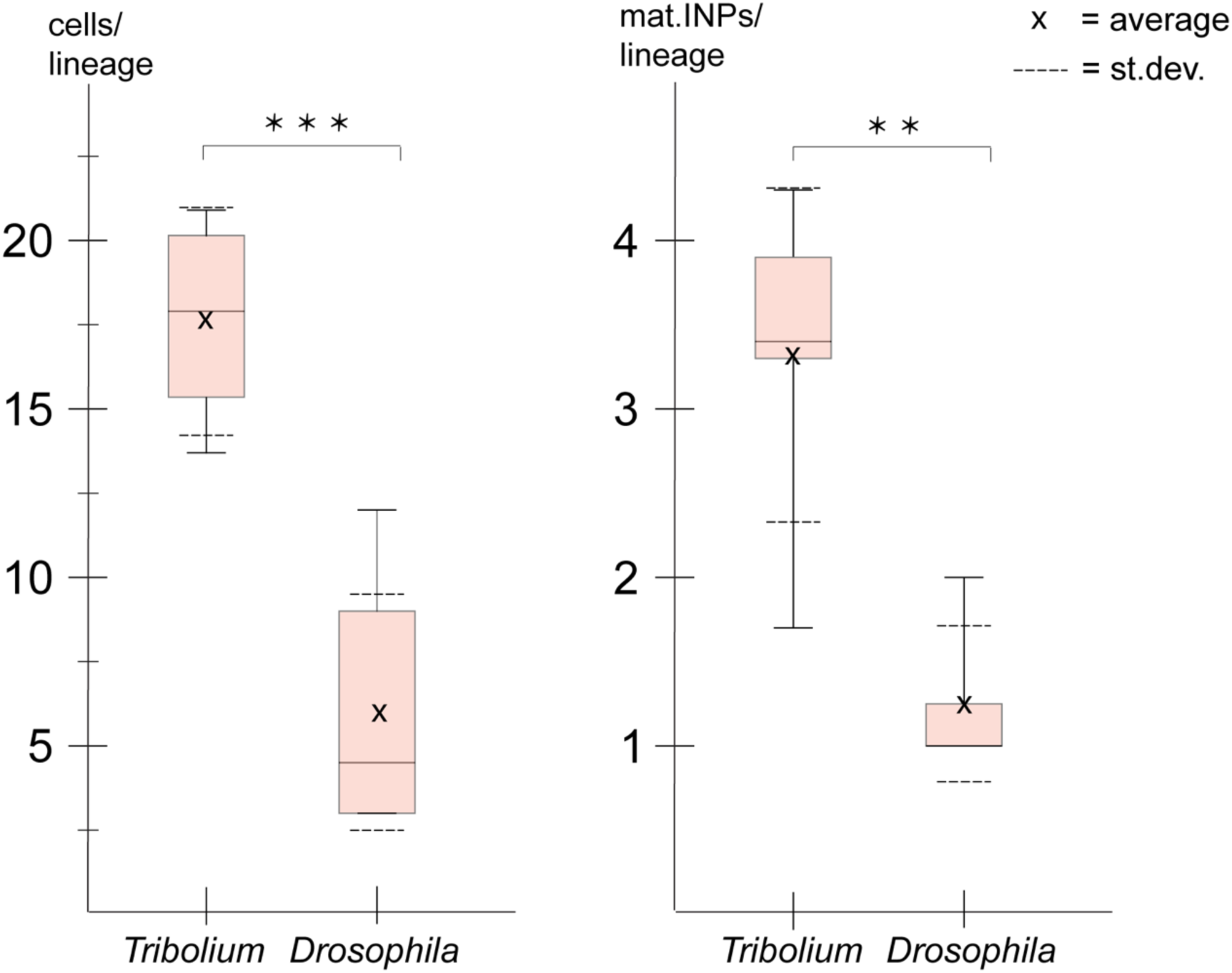
Size comparison of *Tribolium* and *Drosophila* type-II NB lineages. Anteriormedian lineages from *Tribolium* stages 13 and 14 (quantified in this work, see table 1 and 2) compared to *Drosophila* stages 15 and 16 (pooled data from [29]). Left: Lineage sizes comprising type-II NBs, INPs and GMCs (*Tribolium* n=4; *Drosophila* n=8). Right: Number of *Tc-dpn*-positive mature INPs (*Tribolium* n=5; *Drosophila* n=8). T-test p-value significance level (t-test; *** ≘ P < 0.001, ** ≘ P < 0.01).

Due to a lower quality of the image data from the posterior group which is in a deeper layer (see Fig. 3) we were not able to quantify cells of this cluster in *Tribolium*. In *Drosophila,* the posterior lineages are larger (av. 10 cells [29]) than the anterior and median ones, but they are still smaller than the anterior lineages of *Tribolium* at the stages that were investigated. We also found that the embryonic type-II NB lineages in *Tribolium* comprised significantly more mature INPs (*Tc**-**dpn*+/ *Tc**-**fez*+) than the corresponding lineages in *Drosophila* (Fig. 9). The described observations on the higher numbers of progenitor cells in *Tribolium* when compared to *Drosophila* correlate with a previously described earlier (i.e. embryonic) formation of a functional central complex in beetles, which is present in the first larval stage [3].

### Type-II NBs and their lineages are differentially marked by the head patterning transcription factors *Tc-six4* and *Tc-six3* but do not express *Tc-otd*

The anterior-most embryonic insect head is divided into different territories by highly conserved developmental transcription factors that give rise to different parts of the protocerebrum [11]. Hence, we wondered in which of these molecular territories subsets of type-II NB lineages are located, as neuroectoderm regionalization genes are candidates for giving specific spatial identities to individual lineages. We tested co-expression with the previously characterized head patterning transcription factors *Tc-otd*, *Tc-six4*, *Tc-six3*, known to be active in the developing protocerebrum (*Tc-otd*) and in the anterior medial region of the neuroectoderm, where type-II NBs emerge (*Tc-six3*, *Tc-six4*) [11,19,55,56]. We found no co-expression of fez-mm-eGFP marked cells of both the anterior medial and posterior lineages with *Tc-otd*. Rather, *Tc-otd* was expressed in the surrounding embryonic head tissue. *Tc-otd* was also absent from the type-II NBs themselves (Fig. 10 panel A-B). However, *Tc-otd* might be expressed in neural cells derived from these lineages as we can detect expression in the central area of the brain lobes where for instance skh cells are located (Fig. 8 panel A, F; Fig. 10 panel A-B). Interestingly, we found that *Tc-six4* is expressed specifically in that area of the head lobe in which the type-II NBs clusters 1-6 of the anterior medial group first differentiate. At stage NS13 *Tc-six4* marks both, type-II NBs and fez-mm-eGFP expressing INPs and some surrounding cells (Fig. 11, panel A). At the following differentiation stage (NS14) *Tc-six4* still marks the type-II NBs clusters of the anterior medial group but is not expressed in the fez-mm-eGFP INPs (Fig. 11 panel B). It is also not expressed in the posterior group of type-II NBs but only marks type-II NBs of the anterior medial group. Finally, we found that *Tc-six3* marks, and is restricted to, the anterior-most lineages 1-4 including both the type-II NBs and the fez-mm-eGFP marked INPs/GMCs (Fig. 11 panel C). These results reveal candidates for distinguishing type-II identity from type-I NBs in the brain. Only the latter is possibly marked by *Tc-otd* as *Drosophila otd* is expressed in type-I brain neuroblasts [57]. By contrast, *Tc-six4* only marks type-II NBs and appears to be an early determinant of the anterior medial group. *Tc-six4* is also a candidate for distinguishing the anterior medial from the posterior cluster. The results also reveal a subdivision of the anterior cluster by *Tc-six3*, which marks the four most anterior type-II NBs.

**Figure 10.**
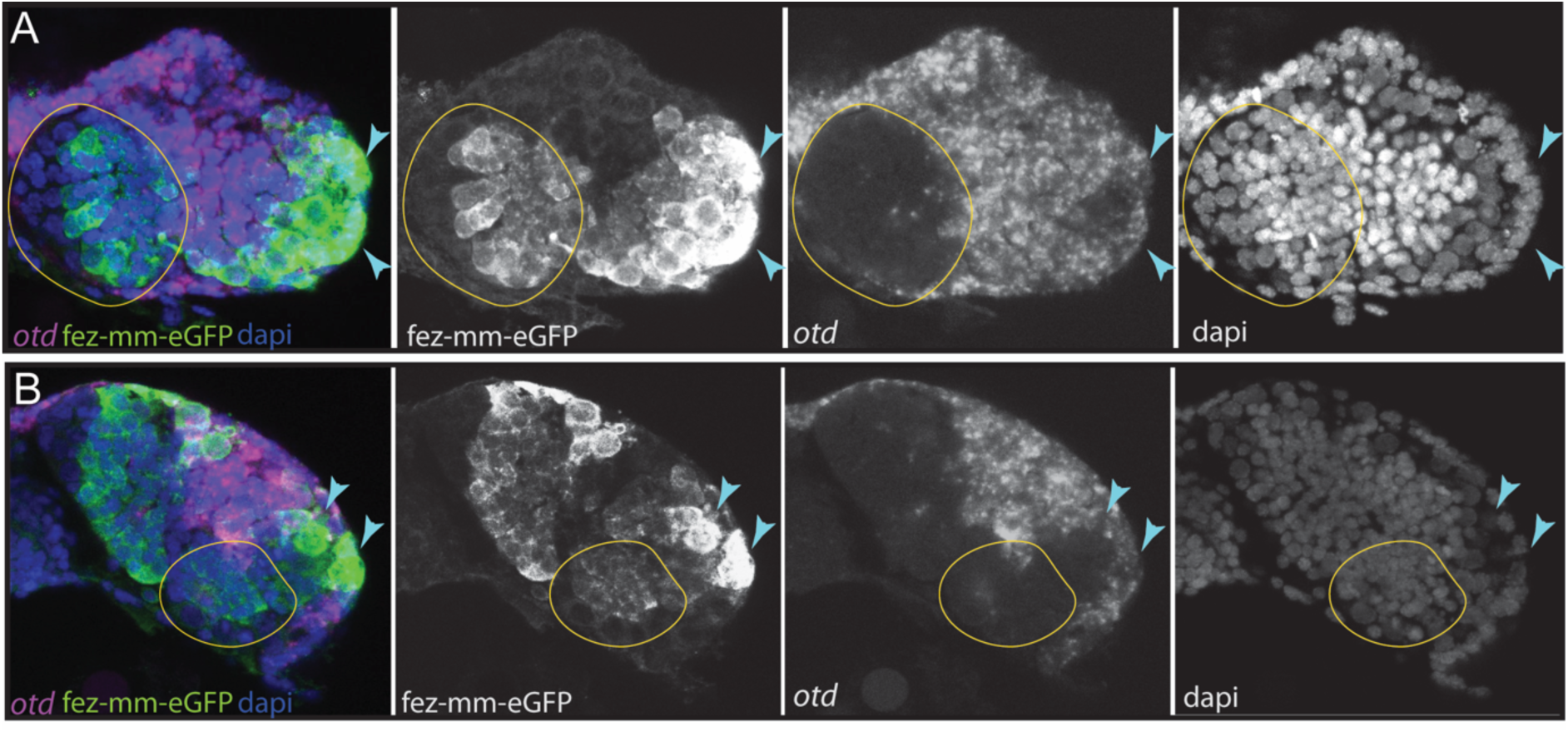
Expression of *Tc-otd* in relation to the fez-mm-eGFP lineages. A/B) RNA *in situ* hybridisation for *Tc-otd* in combination with GFP antibody staining and dapi labelling of nuclei, A and B show projections of different levels of NS14 embryos, right head lobe. **A)** *Tc-otd* is neither expressed in type-II NBs nor the in the fez-mm-eGFP positive INPs of the anterior medial group (both positioned within encircled area) but it is expressed in the directly surrounding tissue. **B)** Likewise, *Tc-otd* is not expressed in type-II NBs or fez-mm-eGFP cells of the posterior group (encircled area). A lateral area of *Tc-fez/erm* and *Tc-otd* co-expression (**A/B**, blue arrow) likely is part of the eye anlagen [19].

**Figure 11.**
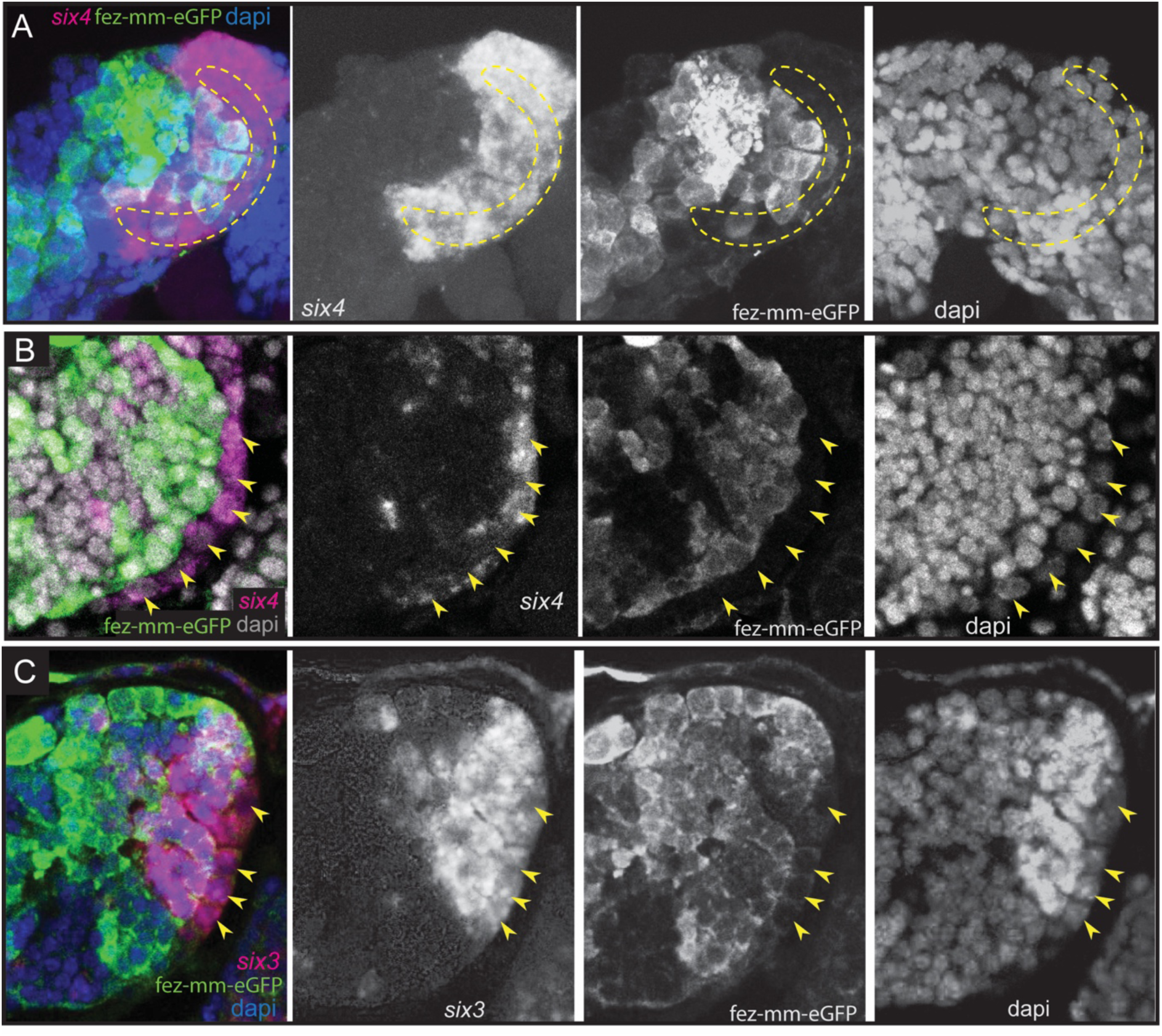
Expression of *Tc-six4* and *Tc-six3* in type-II NBs and INPs. A-C) RNA *in situ* hybridisations of *Tc-six4* and *Tc-six3* in combination with GFP antibody staining reflecting fez-mm-eGFP expression, dapi labelling of nuclei, left head lobes shown. **Panel A)** Stage NS13; *Tc-six4* is expressed in an area encompassing type-II NBs (positioned in encircled area) and mm-eGFP expressing INPs. **Panel B)** Stage NS14; *Tc-six4* is now restricted to type-II NBs and their youngest progeny that do not express fez-mm-eGFP (yellow arrowheads). **Panel C)** Stage NS14; *Tc-six3* specifically marks the four anterior-most lineages. It is expressed in the type-II NBs at the base of the lineages (yellow arrowheads) and in the fez-mm-eGFP positive INPs and possibly also in GMCs. It is absent from the remaining lineages of the anterior medial group and is also not seen in the posterior group (not shown).

## Discussion

### Evolutionary divergence of number and grouping of embryonic type-II NB lineages between beetle, fly and grasshopper

We have identified a total of 9 type-II NB lineages on each side of the *Tribolium* embryonic head. This represents a major evolutionary divergence as in both, the hemimetabolan grasshopper *Schistocerca gregaria* and the fruit fly *Drosophila melanogaster* only 8 embryonic type-II NBs were identified [29,38]. Our finding of an additional type-II NB lineage at the embryonic stage shows that the number of type-II NBs is not fixed across insects and suggests that evolutionary divergence in brain development between species may be driven by differences in the number of progenitor lineages. To test this, further research is required to determine where the additional lineage in *Tribolium* is located and if it is associated with morphological differences of the brain.

Although in the frame of this work we did not perform lineage tracing we can speculate about which NB might be lacking in fly embryos based on our data. In the hemimetabolan *Schistocerca* there is no obvious clustering of type-II NBs, and they are all arranged in one row at the medial rim of the head lobes. In *Tribolium* we have observed an arrangement of the type-II NBs into two groups per side, one large anterior medial group containing seven type-II NB clusters and one posterior group of two type-II NBs. The anterior six of the anterior medial group are arranged in one row like in the grasshopper. In *Drosophila* there are three described groups referred to as the anterior, the middle and the posterior cluster [29]. The posterior cluster consists of two type-II NBs and therefore most likely corresponds to the posterior two type-II NBs in *Tribolium*. The *Drosophila* middle and anterior cluster are most likely equivalent to the anterior medial group of *Tribolium.* However, the type-II NB 7, which we assigned to the anterior medial group, but which is a bit separated and produces offspring into the opposite direction (see Fig. 2 C-II, white arrow) might be the one that does not have a homologue in the fly embryo. The identification of more specific spatial markers for type-II NBs (in addition to *Tc-six3* and *Tc-six4*) or lineage tracing tools are required to identify the neuroblast, which is not present in fly embryos. Subsequently, it would be especially interesting to see what the role of the additional type-II NBs in the beetle is and which neuropile it contributes to.

Summing up, there is a tendency towards grouping of type-II NBs into subsets in the holometabolan models that was not observed in the only hemimetabolan studied so far. A ninth type-II NB was only found in the *Tribolium* embryo represents a striking developmentally divergent feature, but its role and the homologisation of individual type-II NBs between the different insects requires further studies.

### Gene expression identifies homologous cell types and suggests conservation of gene function between fly and beetle type-II NB lineages

The genes defining the different stages of differentiation in type-II NB lineages have been identified and intensively studied in *Drosophila,* but it had remained unclear, in how far the respective patterns and mechanisms were conserved. Our results show a large degree of conservation of expression and probably also function – at least in holometabola. We identified type-II NBs by their larger than average size [40] and the expression of the signature marker *Tc-pnt* adjacent to a group of *Tc-fez/erm* expressing cells (INPs and GMCs). In *Drosophila* both these factors have key functions in defining the developmental potential of type-II NBs and INPs: Pnt suppresses Ase in type-II NBs and promotes the formation of INPs [45]. Loss of *pnt* expression leads to a de-differentiation of INPs as *fez/erm* is no longer repressed in young INPs [44]. Fez/Erm controls proliferation of INPs by activating Pros and prevents dedifferentiation of INPs into type-II NBs [43]. We show that also in *Tribolium* INPs undergo a maturation process, during which expression of *Tc-pnt* stalls and expression of *Tc-fez/erm* is switched on. Despite both factors characterising different stages of INP maturation they are not completely mutual exclusive and we observed a small window of overlap in immature-II INPs, highly suggestive of a conserved role of Pnt in the activation of *fez/erm* [44] (Fig. 12). *Drosophila dpn* is a neural marker [58] expressed in type-II NBs and again in mature INPs, leaving a gap of expression in immature-I and II INPs. *Drosophila ase* is repressed in type-II NBs but expressed together with *fez*/*erm* in INPs and GMCs where its expression overlaps with *pros* [59]. We found identical dynamics of gene expression in *Tribolium* (see Fig. 12) suggesting conservation of gene function within the lineages and a conserved process of INP maturation.

**Figure 12.**
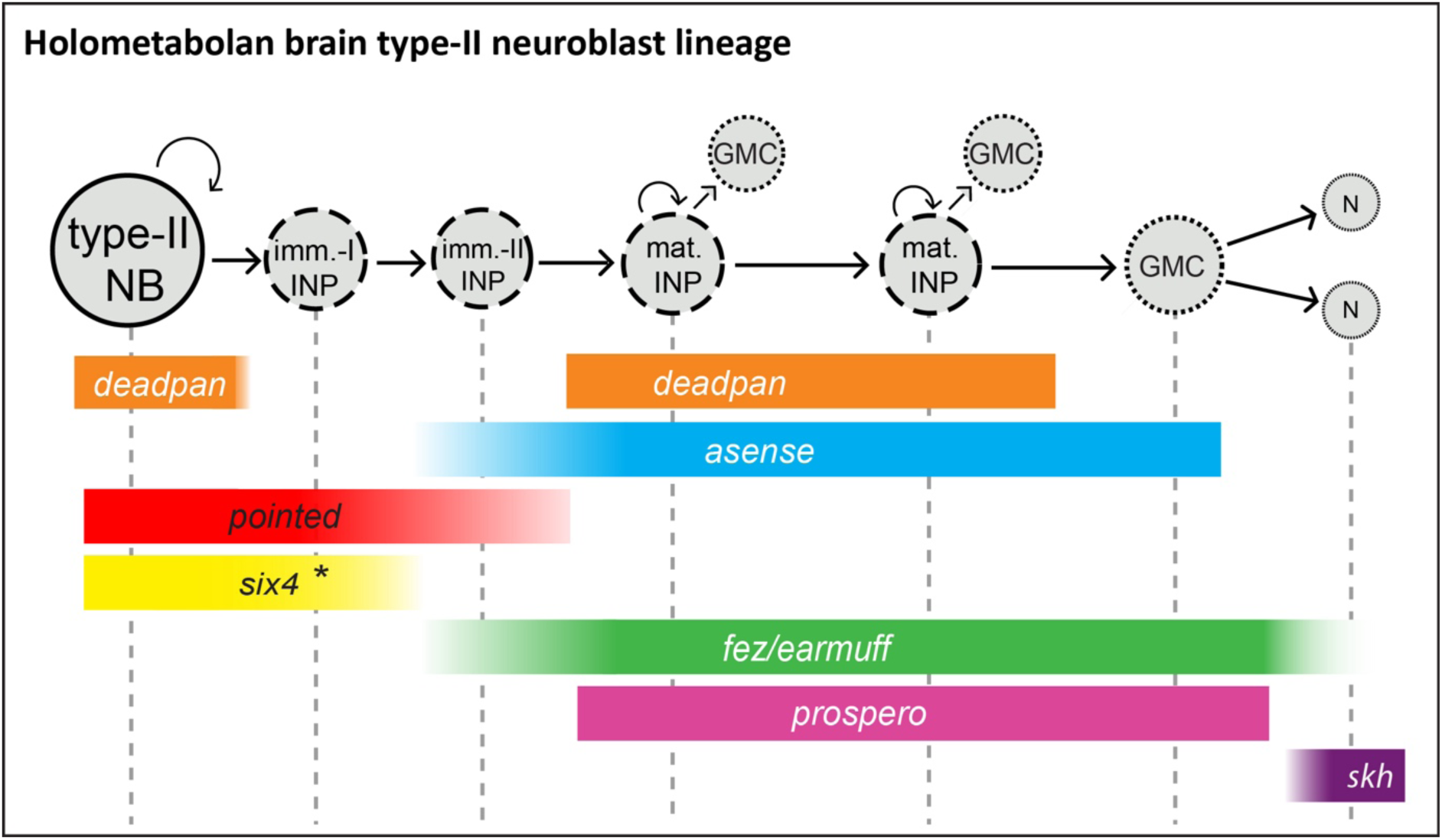
Conserved aspects of gene expression in type-II NB lineages of the holometabolan models *Drosophila* and *Tribolium*. Overview on the different cell types within one lineage (type-II NBs, INPs, GMCs and central complex neurons), their mitotic activity, and summary of the expression of key factors in the respective cell types. **I**NPs are divided into immature-I, immature-II, and mature INPs with each subtype having a unique expression profile. Note that each lineage can have several mature INPs. We strongly assume that glia are also produced from these lineages in *Tribolium*, as they are in *Drosophila* [30]. * In the *Tribolium* embryo *Tc-six4* is only expressed in the anterior medial lineages, and its expression is overlapping with *Tc-fez/erm* at an earlier stage (NS12) whereas it is restricted to type-II NBs and immature-I INPs, mutually exclusive with *Tc-fez/erm*, at a later embryonic stage (NS14). *Drosophila six4* was found in all 8 larval lineages [47]. The scheme is based on this work and [30,41,43,44,47]. NB= neuroblast; imm.-I/-II INP = immature-I/-II intermediate progenitor; mat. INP= mature intermediate progenitor; N=neuron; skh=shaking hands (enhancer trap) [41].

In conclusion, expression of the genes examined in this study in type-II NBs and their lineages is highly conserved in fly and beetle, which is implicit of conserved gene function and a conserved process of central complex formation.

### Divergent timing of type-II NB activity and heterochronic development of the central complex

Previous work described a heterochronic shift in central complex formation between fly and beetle [3,10]. In *Tribolium* parts of the central complex develop during embryogenesis. It becomes functional at the onset of the first larval stage, and is presumably required for the more complex movements of the beetle larvae using legs [3]. In both *Drosophila* and *Tribolium* the anterior groups of type-II NBs (anterior medial group of *Tribolium* and anterior and middle cluster of *Drosophila*) are fully present in the last 3^rd^ of embryogenesis, with the beginning of germ band retraction [8,29,60]. However, in the *Drosophila* embryo only a limited number of INPs and GMCs are produced before they enter a resting stage prior to hatching [29]. By contrast, we show that *Tribolium* type-II NB lineages are much larger at the embryonic stage compared to *Drosophila* [29]. They also include more mature, *Tc-dpn* expressing cycling INPs which have the capability of driving the increase in lineage size. We believe that this increased activity may contribute to the embryonic development of a functional central body and protocerebral bridge [3]. This hypothesis does however require further confirmation by lineage tracing experiments and by testing how manipulating central genes acting within the lineages will affect the timing of central complex development. In addition, differences in cell number in the individual central complex neuropiles between *Tribolium* and *Drosophila* have to be assessed, as the higher number of progenitors may also reflect a higher number of neurons present in the adult beetle fan shaped body or may be due to a slower rate of division and a resulting accumulation of progenitor cells. Lastly, we do not yet have any information whether *Tribolium* type-II NBs, or *Tribolium* neuroblasts in general, also enter a stage of quiescence at the end of embryogenesis as they do in *Drosophila* [29] and what role neurons born in the 1^st^ larva play in *Tribolium*.

Another emerging question is if and when *Tribolium* type-II NB lineages are active during larval development and how that relates to the described steps of central complex development. After preformation in the embryo *Drosophila* type-II NBs have their main period of activity in the 3^rd^ larva and the lineages are much larger and contain more INPs than the *Tribolium* embryonic lineages [30]. Therefore, it would be interesting to see if type-II NB lineages are present and active in the late larval brain of *Tribolium*.

Taken together, we have for the first time identified and molecularly characterized embryonic type-II NBs and INPs as the major central complex progenitor pool in an insect outside *Drosophila*. We have observed quantitative differences within these lineages when compared between *Tribolium* and *Drosophila*. Based on these findings we hypothesize that increased division behaviour of type-II NB lineages is required for the embryonic central complex development as found in beetles. Our findings set the basis for future work on the relationship of the identified type-II NB lineages and the timing of central complex development.

### The placodal marker *Tc-six4* may be responsible for the differentiation and the spatial identity of the anterior medial group of type-II NBs

The different type-II NB lineages contribute to different parts of the brain. For instance, only the anterior four type-II NB lineages form the w, x, y, and z tracts of the central complex in flies and grasshoppers. Hence, there must be signals that make them different despite the conserved and well-studied sequence of neural gene expression and functional interactions in *Drosophila* (Fig. 12) [30,43,44,59]. In the ventral nerve cord, the different identities of the NBs are specified by the combination of spatial patterning genes expressed in the neuroectoderm at the time of delamination [24]. However, little is known about these signals for type-II NBs. In *Tribolium* the domains of head patterning genes in the brain neuroectoderm are well characterized and some highly conserved territories like the anterior-medial *six3*-positive domain were described and suggested to give respective NBs different spatial identities [11,19,55]. The *Tc-six4* expression domain is also largely coincident with an embryonic structure termed insect head placode, a neurogenic, invaginating tissue [56,61] potentially homologous to the vertebrate adenohypophyseal placode which is also marked by *six4* [11,56]. A placodal origin of *Drosophila* type-II NBs has been shown for one type-II NB, and it is assumed that the other type-II NBs also stem from placodal tissue [29,50,62].

*Tc-six4* is also a very interesting factor with regards to the specification of type-II NBs because it is expressed only in the anterior group of type-II NBs. Hence, it could contribute to distinguishing their developmental fate from the posterior group. This is however different in the *Drosophila* larva. Here Six4 is expressed in all eight lineages where it prevents the formation of supernumerary type-II NBs and a premature differentiation of INPs [47]. In the *Tribolium* embryo Six4 may have a similar rate limiting role within the anterior group of type-II NBs. However, its early expression in a small part of the neuroectoderm [56] and the delamination of the anterior-medial type-II NB group within this domain also hints at an instructive role of *Tc-six4* in the formation of the anterior medial group.

### *Tc-six3* marks a subset of type-II NBs whereas *Tc-otd* is absent from all lineages

Importantly, we found that in *Tribolium* late embryogenesis, *Tc-six3* is expressed specifically within the lineages of type-II NBs 1-4 of the anterior medial group, but not in the other linages. In grasshopper and fly these anterior 4 lineages (DM1-4) give rise to the z, y, x and w tracts [25,38] and contribute crucially to the development of protocerebral bridge and central body. These tracts form a major midline crossing neuropile of the central body which serves as a scaffold in the development of this structure [3,25]. We conclude that in *Tribolium* the evolutionary ancient *six3* territory gives rise to the neuropile of the z, y, x and w tracts. This might well be the same in other insects such as flies and grasshoppers but has to our knowledge not been looked at in detail in these models. In the beetle the central body is also missing in weak *Tribolium Tc-six3*-RNAi phenotypes, suggesting that *Tribolium six3* is required for the formation of this structure [19]. Given that central complex neuropile is marked by additional factors such as *Tc-*foxQ2 (*Drosophila* fd102c [63]) and *Tc-rx*- [3,17], it would be interesting to see the relationship of additional spatial patterning genes with regards to the type-II NB lineages in future studies.

The gene *otd* is a marker of the posterior protocerebrum and *six3* and *otd* are believed to subdivide the embryonic anterior brain into two major domains [55]. This subdivision is also part of the recently revisited concept of an ancestral division of the insect protocerebrum into archicerebrum (*six3*-positive) and prosocerebrum (*otd*-positive) [11]. As discussed above, *Tc-six3* is expressed in the anterior-most four type-II NB lineages, but unexpectedly, we found that *Tc-otd* is specifically absent from all type-II NB lineages, suggesting that its inhibition is required for the development of all these lineages. Interestingly, the lineages of type-II NB 5 and 6 (and presumably of type-II NB 7) of the anterior cluster and the posterior type-II NB lineages do neither express *Tc-otd* nor *Tc-six3* showing that there are protocerebral structures expressing none of these conserved markers in development, questioning the assigned generalized role of these two factors in subdividing the protocerebrum [11,55].

In summary, our findings are just the beginning of the quest for the genes required for the identity specification of type-II NB lineages. The genes known to be involved in head patterning are an excellent starting point for this purpose [11,18].

## Methods

### Creation of transgenic reporter line

Using CRISPR-Cas9 we have inserted a transgene into the *Tc-fez/erm* locus that includes *egfp* behind the *Tribolium* basal heat shock promoter (bhsp68), which is not active by itself but can be activated by nearby enhancer elements [64,65]. The transgene also contained the coding sequence of Cre-recombinase (not used in this work) transcribed from the same promoter, and the eye pigmentation gene *Tc-vermilion* behind the eye-specific 3xP3 enhancer element [66] (see Fig. S1 A for map of the transgene). On the repair plasmid the transgene was flanked by CRISPR target sequences derived from *Dm-yellow* and *Dm-ebony*, which were used for *in vivo* linearisation of the transgene.

Using CRISPR Optimal Target Finder [67] we designed three guide RNAs (gRNAs) targeting the genomic region 2.5 kb upstream of the *Tc*-*fez/erm* transcription start site (see table S1). We produced three plasmids where single gRNAs were transcribed from the *Tribolium* pU6b RNA promoter, as described in [12]. We co-injected the three gRNAs plasmids at 125µg/µl each, gRNAs targeting the *Dm-yellow* and *Dm-ebony* sequence at 125µg/µl each, Cas9 helper plasmid (see [12]) at 500 µg/µl and the repair plasmid at 500 µg/µl. Embryos of the white eyed *Tribolium vermilion*-*white* (*vw*) strain (G0) were injected, raised and then bred to *vw*-beetles in individual crosses. The offspring of these crosses (F1) were screened for black eyed heterozygous carriers of the transgene. These were again outcrossed to *vw*-beetles and the heterozygous offspring (F2) were further analysed genetically (see table S2) and with respect to fluorescent signal. Siblings of the F2 generation were then crossed to one another to generate the homozygous line (fez-mm-eGFP). Details of the insertion can be found in Fig. S1 A.

We also used the shaking hands (skh) enhancer trap line (G10011-GFP) which marks central complex cells [41].

### Larval brain dissection, fixation, and staining

Larval brains of the newly generated *Tc-fez/erm* enhancer trap line fez-mm-eGFP were dissected and stained as described in [16]. Brains were mounted in *Vectashield* for microscopic inspection. See table S4 for a list of antibodies and staining reagents used.

### Embryonic RNA *in situ* hybridisation, hybridisation chain reaction and antibody staining

Gene identifiers and primer sequences that were used for amplification can be found in table S3. The fragments were inserted into a pJet1.2 cloning vector. Standard RNA *in situ* probes were synthesised using the 5X Megascript T7 kit (Ambion) according to the manufacturers protocol. Embryo fixation and single colour RNA *in situ* chain reaction in combination with antibody staining to eGFP or phospo-histone H3 staining to mark mitoses was conducted as described in [68]. Embryos were also stained for DNA (dapi) or cell membranes (FM1-43). A full list of antibodies and staining reagents can be found in table S4. Probes for multicolour *hybridisation chain reaction (HCR)* binding *Tc-fez/erm*, *Tc-dpn*, *Tc-ase* and *Tc-pnt* were produced by Molecular Instruments (see table S5). Labelling reactions were performed as described in [69]. Antibody staining to eGFP was performed following the completion of the HCR staining (see [68]). All embryos were mounted in *Vectashield* for microscopic inspection (see table S4).

### Image acquisition and analysis

Multichannel image stacks were recorded using a Zeiss LSM 980 confocal laser scanning microscope. Image stacks consisted of 100-300 slices, depending on the specimen. The resolution ranged from 1024 x 1024 to 2048 x 2048 pixels. Plane thickness was optimized according to width of the pinhole and ranged from 0.1 – 1 µm. We used FIJI [70] for 3 dimensional inspection, to export individual planes or to generate maximum intensity projections. Cropping and adjustment of brightness and contrast was done with GIMP (version 2.10.32) or Adobe Photoshop CS5.

### Evaluation and of cell size and cell number

Cell and nuclear diameters of type-II NBs and a control group were measured based on the image stacks using FIJI [70] across 7 embryos. Lineage size and numbers of mitoses (based on *Tc-pnt* and *fez*-mm-eGFP expression/ anti-PH3 staining) was evaluated by manual counting based on image stacks derived from 4 head lobes of 4 embryos (≘ 28 anterior-medial lineages). The number of *Tc-dpn* expressing intermediate progenitors was evaluated in the same way across 5 embryos (≘ 35 anterior-medial lineages). Reference data on *Drosophila* lineage size (type-II NBs, INPs, GMCs and neurons) and numbers of *dpn*-expressing intermediate INPs were extracted from [29]. Statistics on all numerical data (standard deviations and two-tailed t-tests) were performed using Microsoft Excel 2016.

## Acknowledgements

We would like to thank Claudia Hinners for technical support with molecular biology methods, and Elke Küster for help with screening and beetle stock keeping. We also thank Christoph Viehbahn for valuable feedback on the project. Author Simon Rethemeier was supported by a scholarship of the *Göttingen Promotionskolleg für Medizinstudierende*, funded by the *Jacob-Henle-Programm*.

