## Supplementary material for "Differences in size and number of embryonic type-II neuroblast lineages correlate with divergent timing of central complex development between beetle and fly"

**Table S1:** Target sites for CRISPR-Cas9 mediated *non-homologous end joining* knock-in of the GFP containing transgene (see Fig. S1 A). Guide sequences 1-3 are within 2.6 kb upstream of the transcription start site (TSS).

| CRISPR-guide RNA name | CRISPR-guide RNA sequence incl. PAM |
|---|---|
| ***Tc-fez/erm* upstream 1** | GTGATTACGTGCCGCCGAAG TGG |
| ***Tc-fez/erm* upstream 2** | GCGCTTGCTCGGTTCTCAGT TGG |
| ***Tc-fez/erm* upstream 3** | GCCGTCGTGAGTGAAACGCC AGG |
| ***Dm-ebony*** | GAACCGGGCAGCCCGCCTCC TGG |
| ***Dm-yellow*** | GCGATATAGTTGGAGCCAGC TGG |

**Table S2:** Primer sequences for line fez-mm-GFP insertion site confirmation. See figure S1 for primer binding sites.

| Primer name | Primer sequence | **Distance/ lane in fig. S1 B** |
|---|---|---|
| GFP-5'-rv1 | TGAACTTGTGGCCGTTTACG | 443 bp /1 |
| fez-exon1-rv1 | AACATTAGGTGAGCAGGGCC | |
| **GFP-fw-1** | TTCTTCAAGGACGACGGCAA | 474 bp /2 |
| **P2A-rv** | TCTTCCACGTCTCCTGCTTG | |
| **GFP-fw-1** | see above | 543 bp /3 |
| **Cre-rv1** | GTTGCATCGACCGGTAATGC | |

**A)**

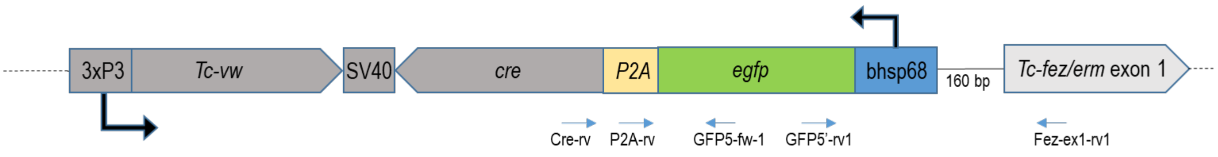


**B)**

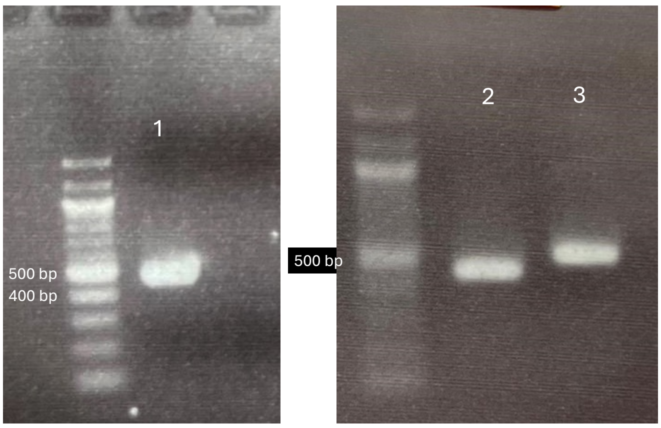

**Figure S1**: **A)** Scheme of inserted construct in fez-mm-GFP line, upstream the *Tc-fez/erm* gene. The orientation of *egfp* is in opposite direction compared to *Tc-fez/erm.* Distance between the bhsp68 promoter and the *Tc-fez/erm* TSS is 160 bp, which corresponds to the distance between CRISPR guide sequence *Tc-fez/erm* upstream 2 (see table S1) and the *Tc-fez/erm* TSS. **B**) Gel electrophoresis with products from PCR using fez-mm-GFP genomic DNA with primers given in table S2 and mapped in figure S1/A).

**Table S3:** Gene and primer sequences used for amplification of fragments for *in situ* probe synthesis.

| Gene | Identifier # | Forward primer | Reverse primer | Fragment length |
|---|---|---|---|---|
| ***Tc-fez/earmuff*** | TC004673 | CAAGCCCTCCATCGTGACCC | GAATCGGAGGCGGAAGTACT | 561 bp |
| ***Tc-pointed*** | TC034783 | GACCGCTTTATTTGCATTGT | TGCTTCTCGTAGTTCATCTTC | 926 bp |
| ***Tc-asense*** | TC008437 | CGTCAGTGTGGTATCCCCTC | GCTGTTCCCACCACTGCATGA | 737 bp |
| ***Tc-deadpan*** | TC005224 | CTCGAGTAACTTACCATTT | CTACCACGGCCTCCACATA | 520 bp |
| ***Tc-prospero*** | TC010596 | ACACAGGGATTCTCGGTCTC | TTGCGTGTCCAAGCAAGAA | 867 bp |
| ***Tc-six4*** | TC003852 | AAGTCGGCGCGAAAGAACGG | CTAAATTTATGGTACTTGAT | 966 bp |
| ***TC-six3*** | TC000361 | ATATGGCGCTCGGACTCGGC | CTCGTACGGTATATATCACG | 1394 bp |
| ***Tc-otd*** | TC003354 | ATGTGGCCTCCAGAGGCAGT | TTAAGCCATATTTGCAAACT | 1116 bp |

**Table S4:** Antibodies and staining reagents.

| Antibody/reagent | Supplier/ reference | Applied concentration (v/v) |
|---|---|---|
| Chicken anti-GFP primary antibody | Abcam/ ab13970 | 1/1000 |
| Rabbit anti phospho-histone 3 primary antibody | Sigma Aldrich/ | 1/100 |
| Anti-chicken Alexa Fluor™ 488 secondary antibody | Life technologies/ A11039 | 1/1000 |
| Anti-rabbit Alexa Fluor™ 647 secondary antibody | Life technologies/ 06-570 | 1/500 |
| Anti-DIG-POD | Roche/ 11633716001 | 1/2000 |
| Anti-Fluoreszein-POD | Roche/ 11426346910 | 1/2000 |
| Tyramide conjugate Alexa Fluor™ 555 | Invitrogen/ B40955 | 1/250 |
| DAPI nuclear dye | Invitrogen/ D1306 | 1/1000 (of 1µg/µl stock sol.) |
| FM1-43 | Invitrogen/ T3163 | 1/1000 (of 5 µg/µl stock sol.) |
| VECTASHIELD® Antifade Mounting Medium | Vector Laboratories/ VEC-H-1000 | As provided by supplier |

**Table S5:** Hairpins, labels and fluorophores used for *hybridisation chain reaction, obtained from Molecular Instruments* (CA, US).

| Probes | label | Fluorophores |
|---|---|---|
| ***Tc-fez/erm*** | B1 | Alexa Fluor 488, 546 |
| ***Tc-deadpan*** | B2 | Alexa Fluor 456, 514 |
| ***Tc-asense*** | B3 | Alexa Fluor 647, 488 |
| ***Tc-pointed*** | B4 | Alexa Fluor 594, 647 |
